# Structure Characterization of Bacterial Microcompartment Shells via X-ray Scattering and Coordinate Modeling: Evidence for adventitious capture of cytoplasmic proteins

**DOI:** 10.1101/2024.11.13.623460

**Authors:** Xiaobing Zuo, Alexander Jussupow, Nina S. Ponomarenko, Nicholas M. Tefft, Neetu Singh Yadav, Kyleigh L. Range, Corie Y. Ralston, Michaela A. TerAvest, Markus Sutter, Cheryl A. Kerfeld, Josh V. Vermaas, Michael Feig, David M. Tiede

## Abstract

Bacterial microcompartments (BMCs) are self-assembling, protein shell structures that are widely investigated across a broad range of biological and abiotic chemistry applications. A central challenge in BMC research is the targeted capture of enzymes during shell assembly. While crystallography and cryo-EM techniques have been successful in determining BMC shell structures, there has been only limited success in visualizing the location of BMC-captured enzyme cargo. Here, we demonstrate the opportunity to use small angle X-ray scattering (SAXS) and pair density distribution function (PDDF) measurements combined with quantitative comparison to coordinate structure models as an approach to characterize BMC shell structures in solution conditions directly relevant to biochemical function. Using this approach, we analyzed BMC shells from *Haliangium ochraceum* that were isolated following expression in *E. coli*. The analysis allowed BMC shell structures and the extent of encapsulated enzyme cargo to be identified. Notably, the results demonstrate that HO-BMC shells adventitiously capture significant amounts of cytoplasmic cargo during assembly in *E. coli*. Our findings highlight the utility of SAXS/PDDF analysis for evaluating BMC architectures and enzyme encapsulation, offering valuable insights for designing BMC shells as platforms for biological and abiotic catalyst capture within confined environments.

## INTRODUCTION

Bacterial microcompartments (BMCs) are organelle-like structures found ubiquitously across microbial phyla that consist of a protein shell encapsulating selected enzyme and cofactor cargo, thus permitting catalytic reaction chemistry to be sequestered from the cytoplasm and tuned by the selective permeability and confined microenvironments of the protein shell.^1-3^ Structures for intact BMCs and their protein shell constituents have been determined with atomic scale resolution from both X-ray crystallography^4^ and cryo-electron microscopy. ^5, 6^ The consistency of protein shell subunit structures when crystallized as isolated proteins compared to those within intact BMC shell architectures point toward the building block assembly mechanism that can be variously combined and tailored to house specific reactions and enables predictive modelling of BMC structures based on subunit composition.^4, 6^ The modularity of shell proteins and their potential for self-assembly in both heterologous expression and in vitro assembly systems present a remarkable opportunity to understand shell protein properties and the principles of self-assembly, as well as to develop a modular tunable framework for spatially confined chemistry that is applicable to both biological and abiotic chemistry. ^7-14^

Several heterologous expression systems have been developed for BMC shell production as stable shells both with and without targeted enzyme cargo encapsulation.^4, 5, 7, 15-17^ While the plasticity of tile-tile building block interactions creates opportunities for mixed-and-matched assembly that can be tuned for selected functional outcomes, it also creates complexity reflected in the pleomorphism of assembled structures.^15^ Similarly, while cryo-EM image sorting algorithms have proven to be of critical value in providing a means to achieve atomic scale structure resolution of BMC shells, there has been less success in using these or other techniques, like dynamic light scattering, SDS-PAGE, or mass spectroscopy, to obtain a quantitative measure of the amount of enzyme cargo encapsulated by BMC shell assembly or to obtain information on the location of the captured cargo with respect to the shell architecture.

In this report, we implement an in-situ structure analysis approach for characterization of a series of BMC shell structures, obtained by heterologous expression of proteins from plasmids carrying the BMC shell genes from myxobacterium *Haliangium ochraceum* in *E. coli*, in solution and under conditions that are directly relevant to biochemical function. In this approach, we calculated both small angle X-ray scattering, SAXS, and corresponding real-space pair density distribution functions, PDDF, from the atom coordinates of BMC shell reference structures and quantitatively compare these to experimentally measured SAXS and PDDF patterns. The analysis was aided by the use of algorithms that permit desktop SAXS and PDDF calculation from atomic coordinates for large, multiple megadalton-scale BMC shell structures.^18^ Comparison of experimental and calculated real-space PDDF curves proved critical for developing and testing of structural models that provide an explanation of variances between solution X-ray experiment and model coordinate structures.

Notably, we found this structural analysis approach allows identification of adventitious protein cargo capture during BMC shell assembly in *E. coli*. In retrospect, while adventitious cargo capture during heterologous expression and assembly could be assumed, there has been no recognition or measurement demonstrating that such capture occurs. In this report, we demonstrate the opportunity to use SAXS/PDDF analyses to quantitate the extent and give information on the localization of adventitious as well as targeted cargo capture during BMC shell assembly. We expect that this approach will prove to be valuable for the development and evaluation of BMC shell architectures for biological and abiotic catalyst capture and as platforms for constructing compartments for catalysis in confinement.

## MATERIALS AND METHODS

### BMC Preparation

We used Gibson cloning to insert C-terminal his-tagged HO BMC-P (Hoch_5814) into a pET11 HO-BMC-H-HO-BMC-T1 co-expression vector (pARH329), described previously, ^19^ to generate the pET11_HT1Phis vector. BL21(DE3) were transformed with pET11_HT1Phis and 2L of LB broth were grown at 37°C until OD600 of 0.8. Protein expression was induced by addition of 0.1 mM IPTG and cells were grown o/n at 22°C. Cells were harvested by centrifugation at 10000xg and stored at -20° C until use. Cell pellets were resuspended in 20ml buffer A (20mM Tris pH 7.4, 50mM NaCl) with 20mM imidazole, supplemented with 200µl 20mg/ml DNAse and lysed by 2 passages through a French Press at 25,000 psi. The lysate was then centrifuged at 38,000xg for 45 min at 4°C and the supernatant was applied on a 5 ml HisTrap HP (Cytiva). The column was then washed with 6cv of wash buffer (20mM Tris pH 7.4, 300mM NaCl, 20mM imidazole) before elution with 2cv buffer A with 300mM imidazole. The elution was then applied on a HR 16/10 MonoQ anion exchange column equilibrated in buffer A and proteins were then eluted with a linear gradient to buffer B (20mM Tris pH 7.4, 1M NaCl). HPhis BMC shells as verified by SDS-PAGE eluted at 34% buffer B and HT1Phis shells eluted at 36% buffer B. Peak fractions were pooled and concentrated and buffer exchanged to Buffer A using Amicon spin filters (30K MWCO).

### SAXS and PDDF Measurements

Small angle X-ray scattering (SAXS) experiments were performed at the 12-ID-B beamline of the Advanced Photon Source, Argonne National Laboratory and beamline 16-ID at the National Synchrotron Light Source II (NSLS-II), Brookhaven National Laboratory. At APS the X-ray wavelength was set to 0.932 Å (13.3 keV), detection used an Eiger2 9M X-ray detector (DECTRIS), and the measurements covered a q range up to 0.85 Å^-1^. At NSLS-II, the X-ray wavelength was set to 0.827 Å (15.0 keV) and a Pilatus3 1M detector (DECTRIS) was used. BMC samples were suspended in 50 mM Tris buffer, pH 7.5 and the protein concentration was adjusted to be in the range of 1-8 mg/ml. To minimize protein damage by X-ray radiation, samples were gently refreshed in a flow cell with a syringe pump.

SAXS patterns, I(Q), were obtained by azimuthally averaging the detector images and binning these as a function of Q = (4*π*/*λ*)sinθ, where 2θ is the scattered angle and *λ* is the X-ray wavelength. I(Q) patterns were obtained for BMC shell samples as the difference between scattering measured for BMC-containing solutions, I_BMC solution_(Q), and the scattering measured from the suspending buffer solution alone, I_buffer_(Q). Pair density distribution functions, PDDF, for BMCs were obtained by indirect Fourier transform of the difference I(Q), using the program GNOM.^20^

### SAXS and PDDF Calculation from Coordinate Structures

SAXS and PDDF patterns from the HO-BMC shell reference structures were calculated using fast SAXS and fast PDDF algorithms in SolX3.^18^ These algorithms allow X-ray scattering calculations for large, multiple megadalton molecular assemblies to be accomplished on desktop computers and to be extended to large Q-ranges.^18^ For example, the algorithms have to been demonstrated to be accurate for simulation of biomolecular wide-angle scattering, (WAXS).^21, 22^ We found the accuracy of these algorithms at high angle to be useful for accurate fitting of SAXS experiments that extended to 0.85 Å^-1^. SolX3 is available for download from https://12idb.xray.aps.anl.gov/solx.html.

To perform SAXS calculations across thousands of BMC shell models, cargo-loaded shells, and molecular dynamics (MD) simulation frames, we used CRYSOL.^23^ SAXS profiles were calculated over the Q-range from 0 to 0.2 Å⁻¹, with 401 data points and 30 spherical harmonics, ensuring small deviation from higher-accuracy tools like SolX3 at this Q-range, while maintaining a high level of computational efficiency, as illustrated in Figure S1. The corresponding PDDF profiles were extracted by evaluating Cα atom pair distances. We note that PDDFs calculated from Cα atoms are essentially indistinguishable from PDDFs calculated from all-atom models, as shown in Figure S2.

BMC shell configurations were assembled using custom Perl scripts that allowed flexible variation in trimer composition (T1, T2, and T3) and the number of pentamers, with a maximum of 12 pentamers per shell and ensuring that the sum of trimers always totaled 20. The script randomly selected trimer and pentamer positions and generated PDB files by merging the components into a complete shell model under the assumption that the overall shell structure remained the same irrespective of the type of trimer and/or missing pentamer tiles. A total of ∼1600 unique shell models were generated to explore how variations in trimer composition and pentamer counts affect the scattering profiles.

To generate protein-filled BMC shells, a second custom Perl script was used to populate the shell interiors with randomly selected E. coli protein structures for the most abundant proteins based on proteomics data (Table S1). Experimental structures with known oligomerization states were used where available. For proteins where experimental structures were not available, we used structures from AlphaFold2 ^24^ as provided in the Uniprot database.^25^ The script ensured that the total weight of the added proteins matched a predefined target, while distributing proteins inside the shell to avoid structural overlap with other the shell or other proteins in the interior. In more detail, selected *E.coli* proteins were initially placed at pre-defined grid positions far enough away from each other to ensure that there was no overlap in the starting positions. A centrosymmetric potential was then applied to guide the proteins to pack within a sphere with a size slightly smaller than the interior diameter of BMC shells during a series of minimization and short molecular dynamics (MD) simulations. The MD simulations were performed using CHARMM with a minimal Cα-based coarse-grained (CG) model to allow close packing without clashes. The interaction potential for the CG model consisted of Cα-Cα harmonic bond terms, Lennard-Jones interactions (ε=0.05 for polar residues, ε=0.2 for hydrophobic residues, σ=3.8Å) and weak electrostatic repulsion for all residues (Q=0.1). The overall structures of folded domains were restrained with a harmonic biasing potential. Optimized CG models were then converted to atomistic models by superimposing the initial atomistic structures to the Cα positions. In this manner, we generated 2400 cargo models for the HT1P and HT1T2T3P shells as well as around 100 models for the HP shell, with varying protein weights and spatial distributions.

### ab initio Molecular Envelope Reconstruction from SAXS Profiles

Three-dimensional (3D) molecular envelopes were reconstructed from SAXS profiles using the ab initio program DAMMIN.^26^ DAMMIN first reads a PDDF file and builds a search space large enough to accommodate the molecular structure under study by taking account of the largest dimension value (Dmax) provided in the PDDF file, and densely fills this volume with small uniform beads, also known as dummy atoms, Figure S3. The program utilizes simulated annealing algorithm to search out an ensemble of beads that their SAXS profile fits the input SAXS data. This ensemble of beads, also often called molecular envelope, represents a structure solution to the SAXS data. If the symmetry of the molecule is known, DAMMIN can apply it to the bead ensemble in the search space which greatly facilitates the molecular envelope search. In this study, an icosahedral symmetry was imposed in the DAMMIN calculations. The resolution of the molecular envelope relies on the size of beads in use, which in turn depends on the selected running mode, and the total volume of search space because there is an upper total bead number limit around 30,000-40,000 for this program.

To achieve an optimal resolution for HT1P reconstruction, DAMMIN calculations were run in Expert mode and used a hollow icosahedron search space with an outer radius of 190 Å, an inner radius of 100 Å, and a bead radius of 5.7 Å. Both experimental SAXS data and the profile calculated from the model structure for HT1P were used for SAXS molecular envelope reconstructions, Figure S3. SAXS data up to Q of 0.154 Å^-1^ were used in the program GNOM^20^ for PDDF and input into DAMMIN for structure reconstruction and fitting. The goodness of the SAXS data fittings in DAMMIN calculations is partially limited by the bead size accessible for this study.

### Molecular Dynamics Simulations

To test the hypothesis that the discrepancy between experimental and computed SAXS profiles were due to nanoscale dynamics of the BMC shells, we conducted molecular dynamics simulations starting from the 6mzx cryo-EM structure. As some residues were missing from the experimental structure, AlphaFold structures from Uniprot ^27^ were taken to fill in the gaps and were aligned to the original cryo-EM structure to generate a complete shell model. The protein model was solvated in a 456 Å cubic box, neutralized, and ionized with 0.15M NaCl in VMD, ^28^ prior to equilibration and simulation with NAMD 3.0.^29^ Noting that water fluxes were imbalanced across the shell, additional water molecules were placed within the shell to balance the volume. Production simulations were carried out for 100 ns to sample structural heterogeneity at short timescales for a thermalized structure. The simulation was performed at 298 K and 1 atm maintained by a Langevin thermostat^30^ with a friction coefficient of 1 ps^-1^ and an isotropic Langevin barostat^31^ with 2 fs timesteps enabled through the use of the SETTLE algorithm.^32^ Long range electrostatics were captured through the particle mesh Ewald method ^33^ with a 1.2 Å grid resolution. The trajectory was analyzed for water flux, the root mean square deviation (RMSD) of the protein components. Snapshots were evaluated to calculate SAXS profiles and to measure their variation over time.

### Proteomic analysis

Proteomic analysis was conducted at the MSU Proteomics Core Facility as described previously.^34^

## RESULTS

### BMC Shell Reference Structures

Figure 1 shows BMC shell reference structures used in the analysis of experimental X-ray scattering for HT1T2T3P, HT1P, HP BMC shells from *Haliangim ochraceum* (HO) that were isolated following heterologous gene expression in *E. coli*. Five structural subunits are used in assembly of HO BMCs: BMC-H, BMC-T1, BMC-T2, BMC-T3, and BMC-P, referred to as H, T1, T2, T3, and P, respectively. H is a hexamer that forms a hexagonal tile, while T1, T2 and T3 are trimers with a duplication in protein sequence and structure that allows them to form similar hexagonal tiles. T2 and T3 each dimerize to form stacked tiles while T1 remains unstacked. P is a pentamer that forms the vertices of the icosahedron.^15^ By controlling which HO BMC subunits are present during expression, several distinct HO shell types, including a full HT1T2T3P and minimal HT1P shells.^15^ Models for the HT1T2T3P shell have been determined from both X-ray crystallography^4^ and cryoEM.^6^ Coordinates from the latter, 6mzx, determined for a HT2P shell, were used in the analysis. A model structure for HT1P shell was obtained by substituting the T1-hexamer 6n06 ^6^ into T2 positions in the HT2P shell structure. Smaller HP shells without trimers are also found to be assembled during full and minimal shell expression and can be isolated separately. A structure has not yet been determined for the HO-HP shell. However, HP BMC shell structures have been determined in other organisms, including the T=3 (6owf) and T=4 (6owg) icosahedral shells from *Halothece sp.*,^35^ shown in Figure 1.

**Figure 1.**
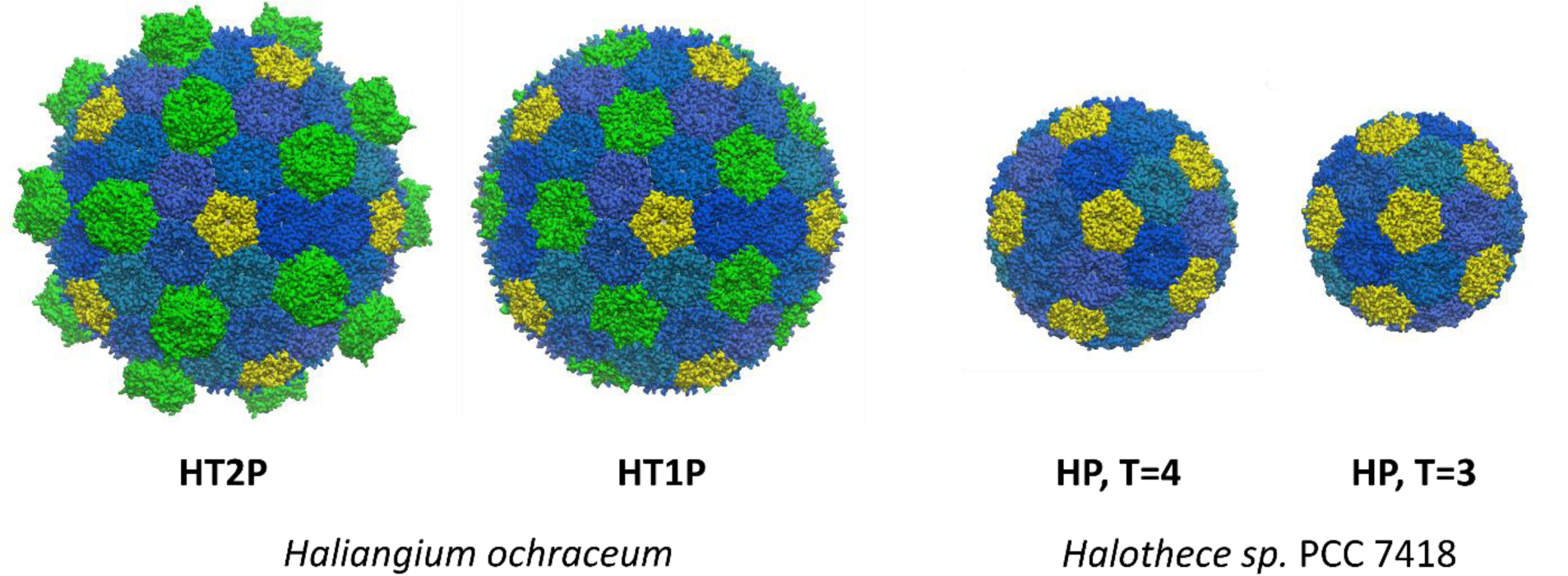
BMC shell reference structures determined by cryo-EM. HT2P (6mzx) and model HT1P shells from *Haliangim ochraceum*, HP T=4 (6owg) and T=3 (6owf) shells from *Halothece sp*. PCCC 7418. Hexamer subunits are shown in blue, trimers in green, and pentamers in yellow.

### SAXS and PDDF Measurements

Experimental SAXS data for HO-HT1P and HO-HT1T2T3P shells are shown in Figures 2A and 2B, respectively. The experimental traces are scaled and plotted with the scattering patterns calculated from the atomic coordinates for the corresponding HO-BMC shell reference structures. Overall, the experimental and calculated scattering patterns show good correspondence. In the small angle region, Q < 0.005 Å^-1^, the scattering curves asymptotically approach the limiting value, I(0), a parameter proportional to the squared molecular mass of the shell. I(0) and the radius of gyration, R_g_, can be extracted from Guinier plots, ^36-38^ log[I(Q)] versus Q^2^. Guinier plots for the HT1P and HT1T2T3P shells, Figures S4A and S4B, respectively, show that the experimentally determined R_g_ for these shells, 118 (±3) Å and 119 (±3) Å, are slightly larger than those calculated from the corresponding reference structures, 108.7 Å and 111.3 Å, respectively. The linear Guinier plots for the experimental HT1P and HT1T2T3P data show that these shell preparations are homogeneous in particle size, with no indication of deviations that would be characteristic of significant interparticle interactions or aggregation.

**Figure 2.**
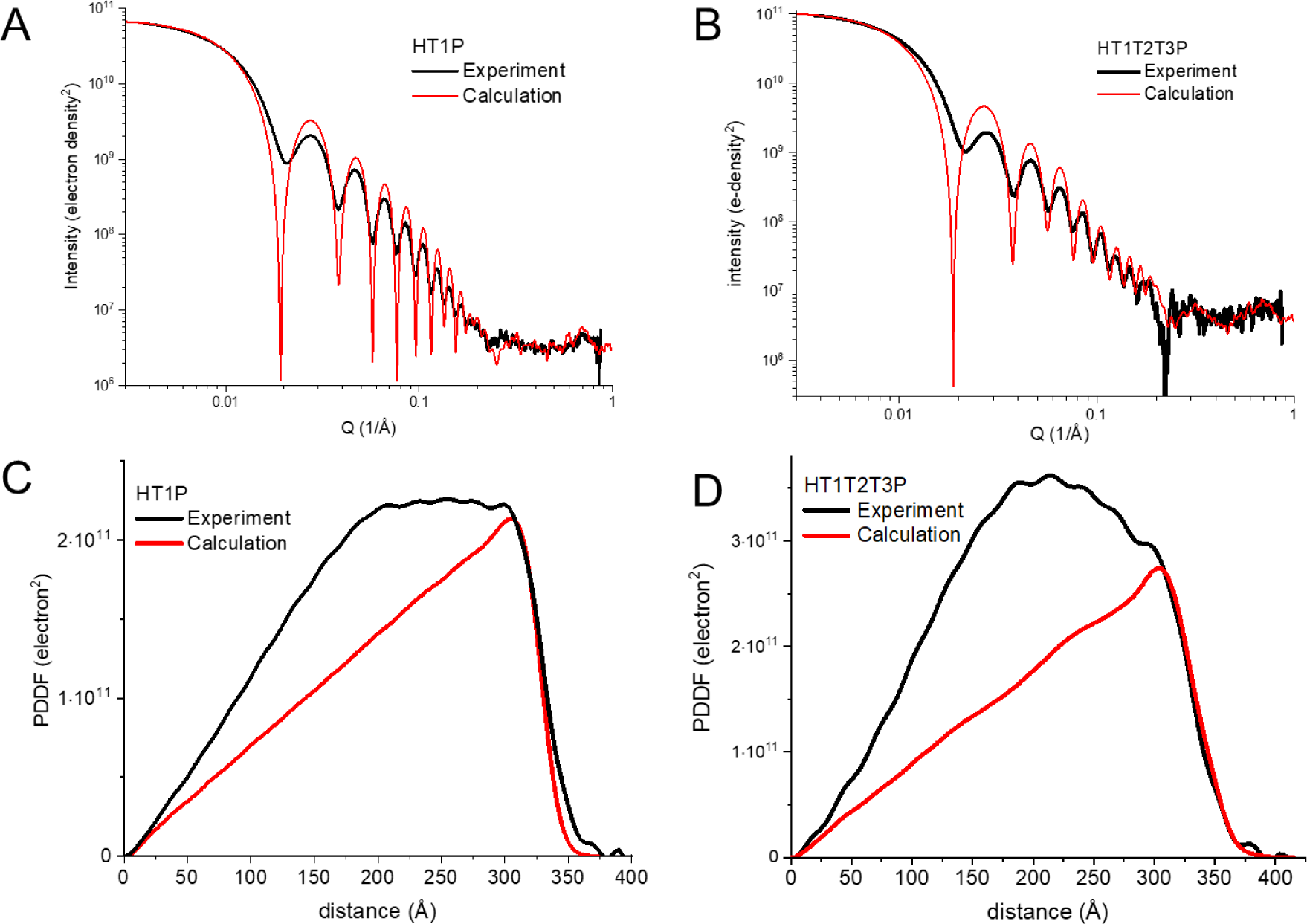
Comparison of experimental and calculated X-ray scattering and PDDF for HT1P and HT1T2T3P BMC shells. **Parts A** and **B**, show experimental (red) and calculated (black) SAXS patterns for HTP and HT1T2T3P shells, respectively. Scattering for the HT1T2T3P shell was calculated directly from the cryoEM atomic coordinate model PDB entry 6MZX, while scattering for the HTP shell was calculated from a coordinate model built from the 6MZX structure, where the HT1T2T3 fragments were replaced by the HT1 partial structure, PDB entry 6N06. **Parts C** and **D** show PDDF patterns corresponding to the scattering shown in Parts A and B. PDDF from experimental scattering data were obtained from indirect Fourier transform using the program GNOM.^26^ PDDF were calculated from atomic coordinates using the program SolX3.^18^

At higher angle, the SAXS patterns for BMC shells show a series of intense inference fringes that are characteristic of approximately spherical, polyhedral structures. Calculations using analytical expressions for core-shell structures, Figure S5 and discussed further below, show that the positions and intensity pattern of the interference fringes are highly sensitive to the diameter, wall thickness, and electron density of the interior space. Apart from the first interference feature, the interference fringes measured for the HT1P and HT1T2T3P shells show close correspondences to those calculated from the reference structures, suggesting the appropriateness of the reference structures as models for the solution state structures. However, the experimental measurement for both HT1P and HT1T2T3P shells show deviations in the shape and position of the first interference feature, seen in the experimental traces by the minimum near Q = 0.2 Å^-1^, and by the dampening of the subsequent oscillatory features. In this context, it is interesting to point out that the higher order interference peaks calculated from the HT2P shell, Figure 2B, are seen to be more strongly damped compared to those calculated from the HT1P coordinate structure, Figure 2A. Presumably, this can be understood to arise from the variation in shell wall thickness produced by the exterior-facing T2 subunits.

Figure 3A shows experimental X-ray scattering data for HO-HP shells compared to scattering calculated from coordinate structures determined for the T=4 (6owg) and T=3 (6owf) HP shells from *Halothece sp.* PCCC 7418.^35^ The prominent interference frequency mismatch with the *Halothece* T=3 HP shell shows that this model structure is much smaller than the HO-HP shell. However, scattering calculated from the T=4 HP structure shows good correspondence with the scattering measured for HO-HP shells, albeit with additional broadening present in the in-situ experiment. This shows that the overall shape and dimensions of the HO-HP and *Halothece* T=4 HP shell are comparable. However, experimental scattering for the HO-HP sample shows deviation from the *Halothece* T=4 HP model in the SAXS region Q < 0.01 Å^-1^. Guinier plots, Figure S6, show that data for the HO-HP shell deviates from the linear plot obtained from the scattering calculated from the *Halothece* T=4 HP structure. The results are indicative of some form of heterogeneity, aggregation or inter-particle interactions for the HO-HP shell preparation. Further, it is interesting to note that in the experimental HO-HP data and calculated scattering from the *Halothece* models, interference features are seen to extend to at least Q = 0.8 Å^-1^, Figure S7, suggesting that it would be possible to use wide angle scattering data with spatial resolution better than 7.85 Å to guide refinement of coordinate models to fit the HO-HP experimental data.

**Figure 3.**
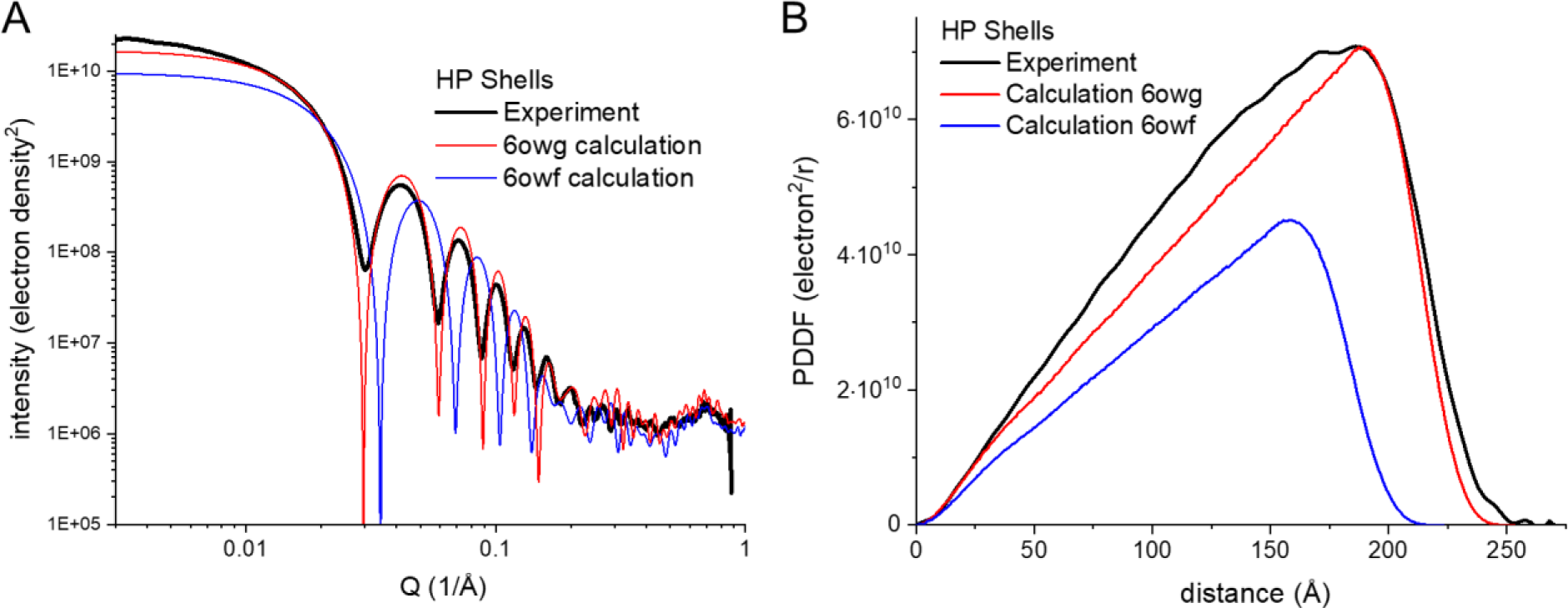
Comparison of experimental and calculated X-ray scattering and PDDF for HP BMC shells. Parts A shows experimental scattering for HO-HP shells (red) compared to a scattering pattern calculated for the T=3 and T=4 HP shells from Halothece sp. PCC 7418, 6owf plotted in blue and 6owg plotted in red, respectively. Part B shows PDDF patterns corresponding to the experimental and calculated scattering curves shown in Part A. The PDDF from atomic coordinates, 6owf and 6owg, give distance maxima of 223 Å and 253 Å, respectively.

In order to gain further insight into structural differences between in-situ BMC shells and reference structures based on variances in SAXS patterns, it useful to compare experiments with model structures based on real space PDDF. For example, PDDF patterns for HT1P and HT1T2T3P shells are shown in Figures 2C and 2D, obtained by indirect Fourier transform of the experiment SAXS data in the Q-range up to 0.30 Å^-1^. Experimental PDDF patterns are plotted with those calculated from the corresponding reference structures. Experimental and calculated PDDF patterns for HP shells are shown in Figure 3B.

PDDF patterns calculated for each of the BMC shell reference structures show a similar triangular shape that scales with the dimension of the shell. An overlap of the HP, HT1P, and H2P calculated PDDF patterns is shown in Figure S8. The triangular shapes can be understood to arise from electron density pairs starting from nearest neighbors and extends across progressively longer chords of the shell until a maximum is reached along the shell diameter at the farthest circumference that maintains the average electron density of the shell. Beyond this the PDDF fall steeply, reflecting “roughness” of the shell outer edge, due to both the shape of the shell and extensions of the amino acid side chains. The longest atom pair distances for the T=3 HP, T=4 HP, HT1P, and HT1T2T3P reference models are 223 Å, 253 Å, 374 Å and 416 Å, respectively. The presence of T2/T3 proteins protruding on the exterior surface of the HT1T2T3P shell can be seen to add undulations to the triangular PDDF shape at distances below the diameter of the shell, and then to extend the distance maximum. Figure S9 shows a comparison of the scaled PDDF for HT1P and HT2P reference structures, illustrating the difference in the long-distance edges. The comparison of PDDF for HT1P and HT2P structures is of interest since it illustrates how protein attachment to the exterior surface of BMC shells would be recognized. The models show that the additional T2 protein protrusions are tracked in PDDF by added pair-distance electron densities within the diameter of the shell and an extension of the long-distance edge of the PDDF, while contributing relatively less in the distance range below the shell diameter.

Figures 2C, 2D, 3B show that the PDDFs measured for HO-BMC shells by experiment deviate significantly from those calculated from reference structures. Characteristic features of these deviations are prominent addition of pair electron density at distances shorter than the diameter of the shells and only relatively minor deviations in long-distance edges of the PDDF. The extent of the added pair electron density between experiment and calculation varies among the different shell types, with the HT1T2T3P shell model having the highest added density pairs within the diameter of the shell, HP the lowest. Further, we have observed that the PDDF patten measured for a single shell type can vary significantly with different preparations. For example, Figure S10 shows SAXS and PDDF measurements made for different HT1P preparations. The PDDF patterns are seen to vary mainly in the amount of added density pairs at distances shorter than the shell diameter.

To further understand the X-ray scattering profiles of BMC shells, calculations have been performed using analytical expressions for hollow spheres and icosahedrons which are two closely related geometric models for BMC shells.^39^ As demonstrated in Figure 4, the scattering profile for a solid sphere (SSp) exhibits Bessel function like oscillation shape with one interference frequency, which is a function of the sphere radius. However, hollow spheres (HSp) have two beat frequencies in the scattering profiles. The fast beat frequency is close to that of the solid sphere with the same outer radius, and moderately low q shifted by the inner radius value. The slow beat frequency of the hollow sphere is dominated by the wall thickness. As shown in both the normal I(Q)-vs-Q scattering profiles and the Q^4^*I(Q)-vs-Q Porod-Debye plot, the slow frequencies are the same for the three hollow spheres with same wall thickness but different inner or outer radii. The slow Q periodicity is a simple function of the wall thickness, i.e., Q periodicity = 2π/thickness.

**Figure 4.**
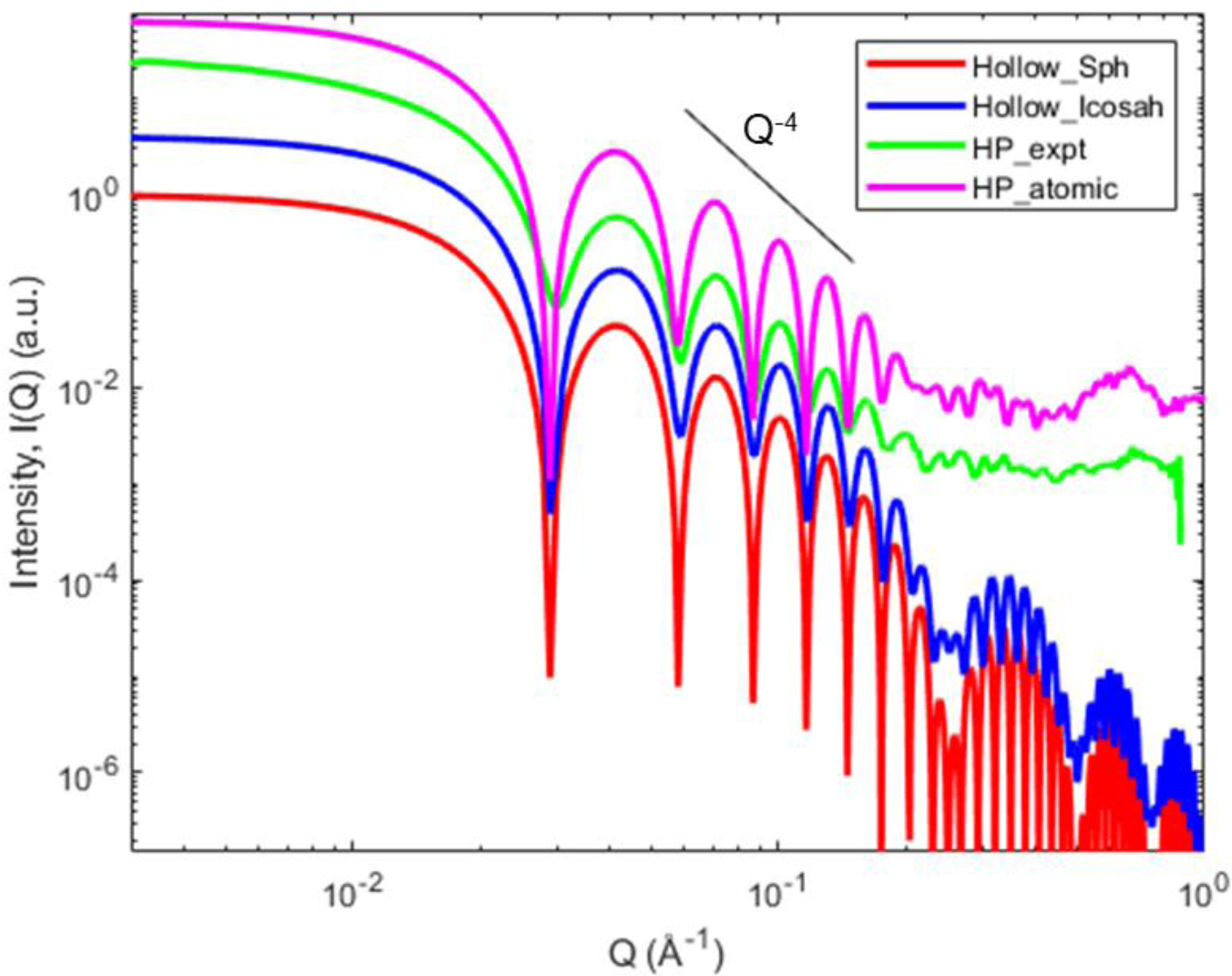
Experimental (green) and simulated X-ray scattering profiles for HP BMC shell and models. Red: calculated scattering profile with a hollow sphere model with a shell wall thickness of 25 Å and an inner hollow core radius of 95 Å. The scattering intensity decays along with q following I(q) ∼ q^-4^ fashion, in the similar slope as shown in Figure S5A. Blue: calculated scattering profile for a hollow icosahedron model with a shell wall thickness of 31.5 Å and an inner radius of 110 Å. Green: experimental X-ray scattering for HP shell. magenta: calculated scattering profile with pdb 6QN1. The profiles were vertically offset for clarity.

As displayed in Figure 4, SAXS calculated for a hollow icosahedron (HIc)^39^ produces a profile (in blue) with two beat frequencies that matches those for a HSp (in red). Again, the slow beat frequency is dominated by the shell wall thickness. Although very similar, there are still a couple characteristic differences in the scattering profiles between HSp and HIc. First, the oscillations of HIc are significantly dampened compared to those of HSp. Second, the oscillation minima of HIc exhibit a different pattern, particularly, the second low Q minimum is shallower than its neighbors. This feature is also seen in the magenta scattering profile that is calculated from atomic structures with icosahedron symmetry, Figure 4.

The fast beat frequency of the scattering profile calculated using atomic coordinates for the T=4 HP shell (magenta trace) match closely to those calculate from the HSp and HIc models in the Q region below 0.25 Å^-1^, indicating these structures are close in the overall dimension and shell wall thickness. The slow beat frequency becomes less pronounced in the high Q region of the atomic simulated profile. The oscillation pattern in the range Q = 0.25 - 0.50 Å^-1^ is complicated by the radii and wall thickness of the atomic assembly, the shape irregularity, and a high flat background. The flat background intensity presumably arises from the electron density fluctuation within the assembly, otherwise the high-Q intensities would decrease roughly in the Q^-4^ fashion as the uniform geometric models. The experimental X-ray scattering profile resembles the calculated one from atomic structure, but the oscillations are further dampened, and the first oscillation shifts slightly towards the higher Q values.

To understand the discrepancy between the experimental and atomic structure simulated X-ray scattering data, 3D ab initio molecular envelopes were reconstructed using program DAMMIN and experimental and simulated data for BMC shell HT1P. In order to facilitate the calculation, icosahedron symmetry was applied. As shown in Figure 5A, DAMMIN successfully produced the empty hollow core-shell structure (in cyan) using the SAXS profile calculated from the HT1P coordinates as an input, as seen by the overlap with the atomic model (in red). In contrast, the molecular envelope reconstructed using experimental HT1P SAXS data yielded a shell structure (magenta structure) having a significant amount of additional molecular mass inside the shell, associated both with the shell wall and interior space. Comparison of the SAXS patterns associated with the ab initio models Figure 5B, shows that the shift in the first interference minima and dampening of the subsequent oscillatory features can be understood to be correlated to the presence of molecular mass inside the BMC shell structures. However, comparison of the DAMMIN fits to input calculated and experimental SAXS patterns, Figure 5B, shows that the discrepancies remaining between input and fitted patterns are likely, at least in part, to be due to the accessible bead size and structure resolution limits in the program.

**Figure 5.**
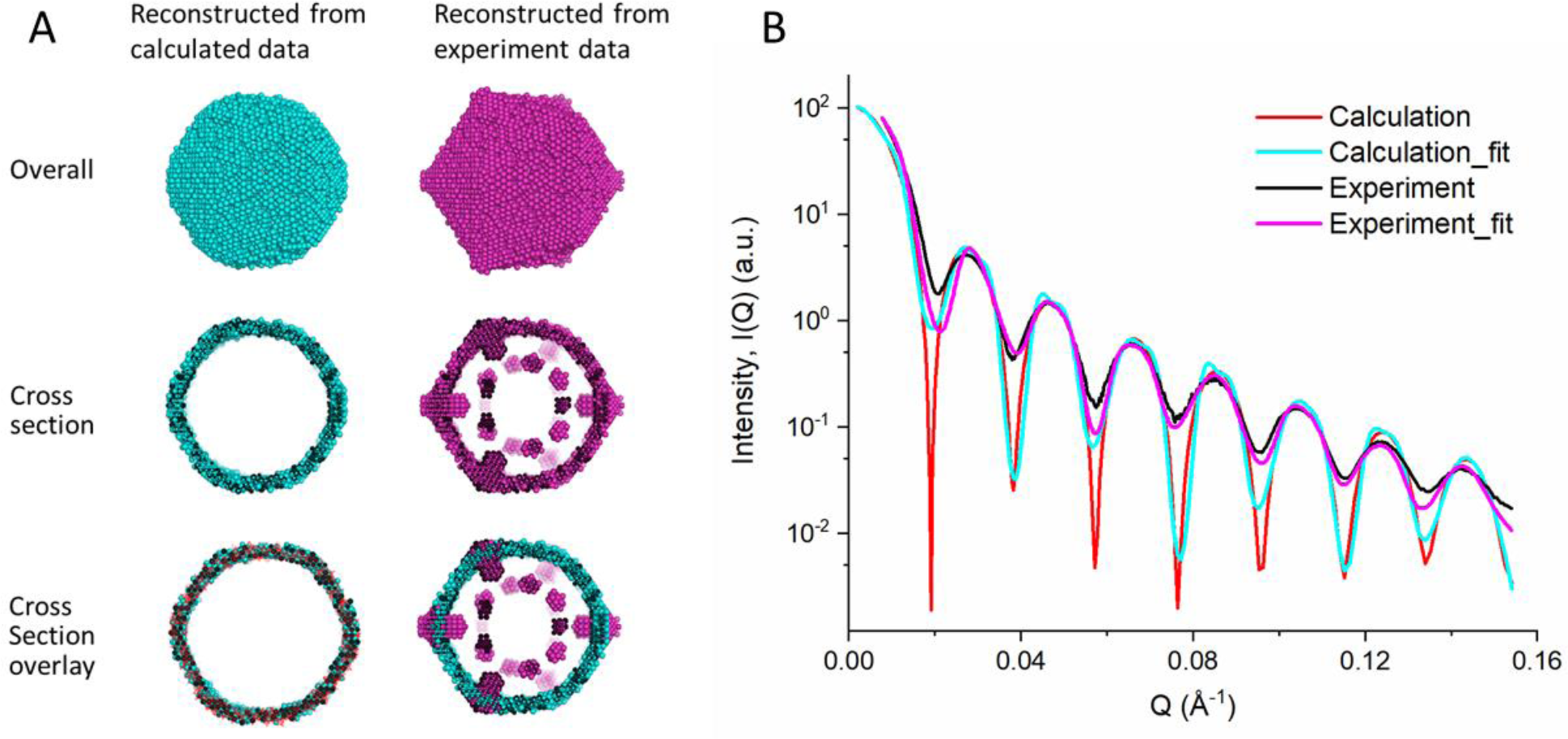
Part A. ab initio 3D HT1P molecular envelopes reconstructed using program DAMMIN from SAXS data sets and comparison among structures. Icosahedron symmetry was applied in SAXS molecular envelope reconstruction, and more details can be found in Materials and Methods section. The molecular envelopes in magenta and cyan were reconstructed from experimental and calculated HT1P SAXS profiles in Fig 2A, respectively. The first and second rows are the overall and cross-section views of the SAXS molecular envelopes. The third row are the cross-section overlays: left, the cyan molecular envelope and atomic structure (in red, PDB entry 6N06); right, the two SAXS molecular envelopes. Part B. SAXS data profiles and DAMMIN fitting in the ab initio reconstruction. Black and Red: HT1P experimental and calculated profiles, respectively, taken from Fig 2A with Q up to 0.154 Å^-1^ and rescaled. Cyan and magenta are DAMMIN fittings, corresponding to the structures in Fig 5A in the same colors.

### Coordinate-based Modeling

While experimentally resolved structures of individual BMC shells provide a strong foundation for understanding their properties, significant discrepancies remain between predicted and experimental SAXS and PDDF profiles. To address this, we generated over 5000 different BMC models, systematically varying several factors that could account for the observed differences. Specifically, we examined the effect of changes in the shell structure by altering the trimer composition and the number of pentamers on calculated SAXS and PDDF patterns, Figures 6A and 6D, and 6B and 6E, respectively. Additionally, we tested the impact of local dynamics through additional molecular dynamics (MD) simulation with the HO BMC shell, Figures 6C and 6F, as well as the effect of mixing different shell types, Figure 7. We note that the possibility of having a mixture of different shell types can be ruled out because of the linearity of Guinier plots, Figure S4, as discussed above, and because the shell preparations used for structural analysis by cryo-EM have been shown to be homogeneous.^6^ Lastly, we tested the effect of encapsulated cargo proteins, Figures 8 and 9.

**Figure 6.**
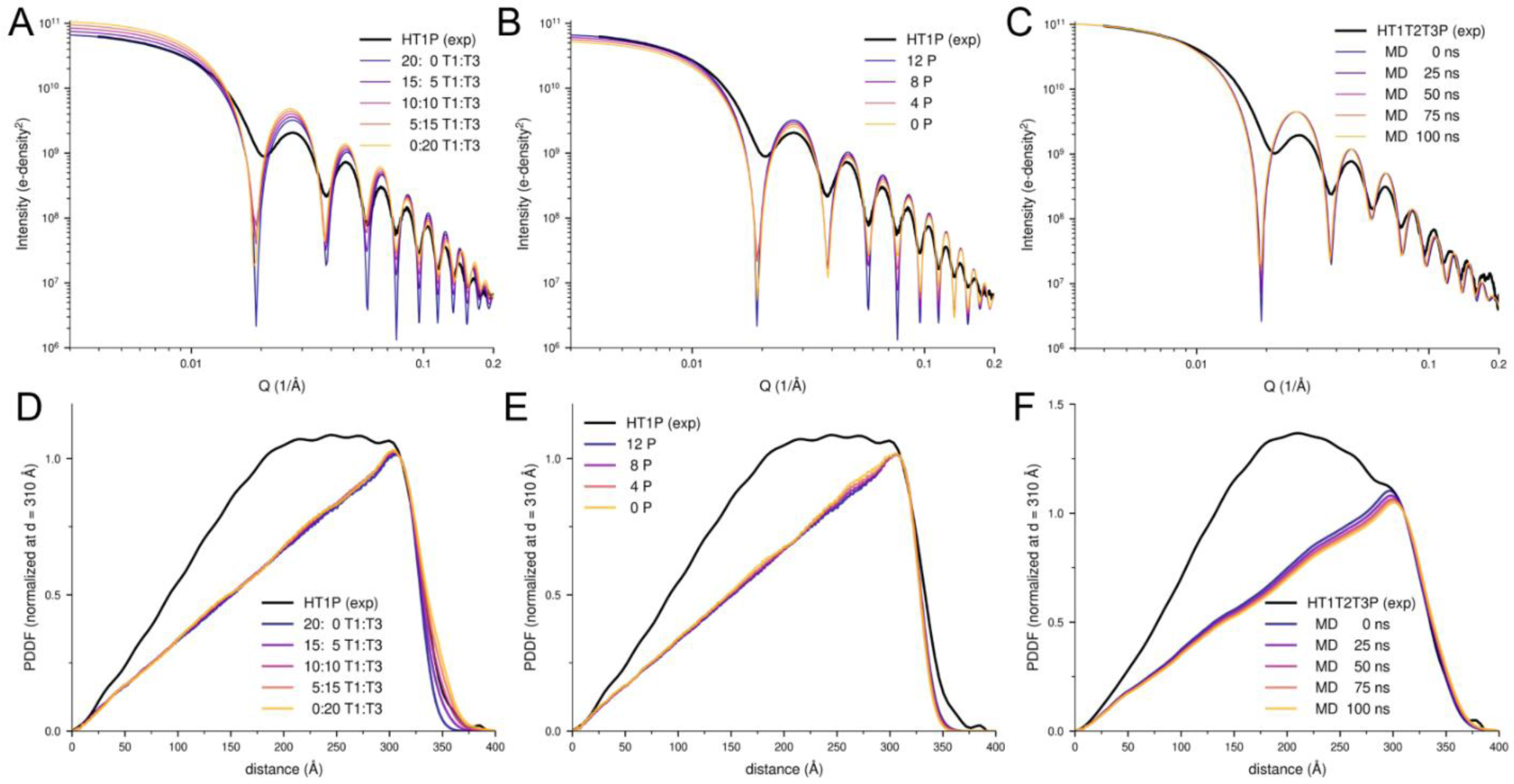
Sensitivity of SAXS and PDDF profiles to structural changes and local dynamics. Parts A to C compare experimental (black) with calculated (colored) SAXS profiles. (A) illustrates the effect of replacing a fraction of T1 trimers with T3 trimers in the HT1P shell. (B) highlights the impact of varying pentamer content on the SAXS profile of the HT1P shell. (C) shows the changes of the SAXS profile over a 200 ns molecular dynamics (MD) simulation of the HT1T2T3P shell. Parts D to F show the corresponding PDDF profiles, normalized at 310 Å. (D) displays PDDF profiles for varying trimer ratios, (E) reflects the influence of pentamer content, and (F) shows the changes in the PDDF profiles during the MD simulation. Overall, local structural conformational changes have a limited impact on the SAXS and PDDF and are not sufficient to explain the differences with the experimental data.

**Figure 7.**
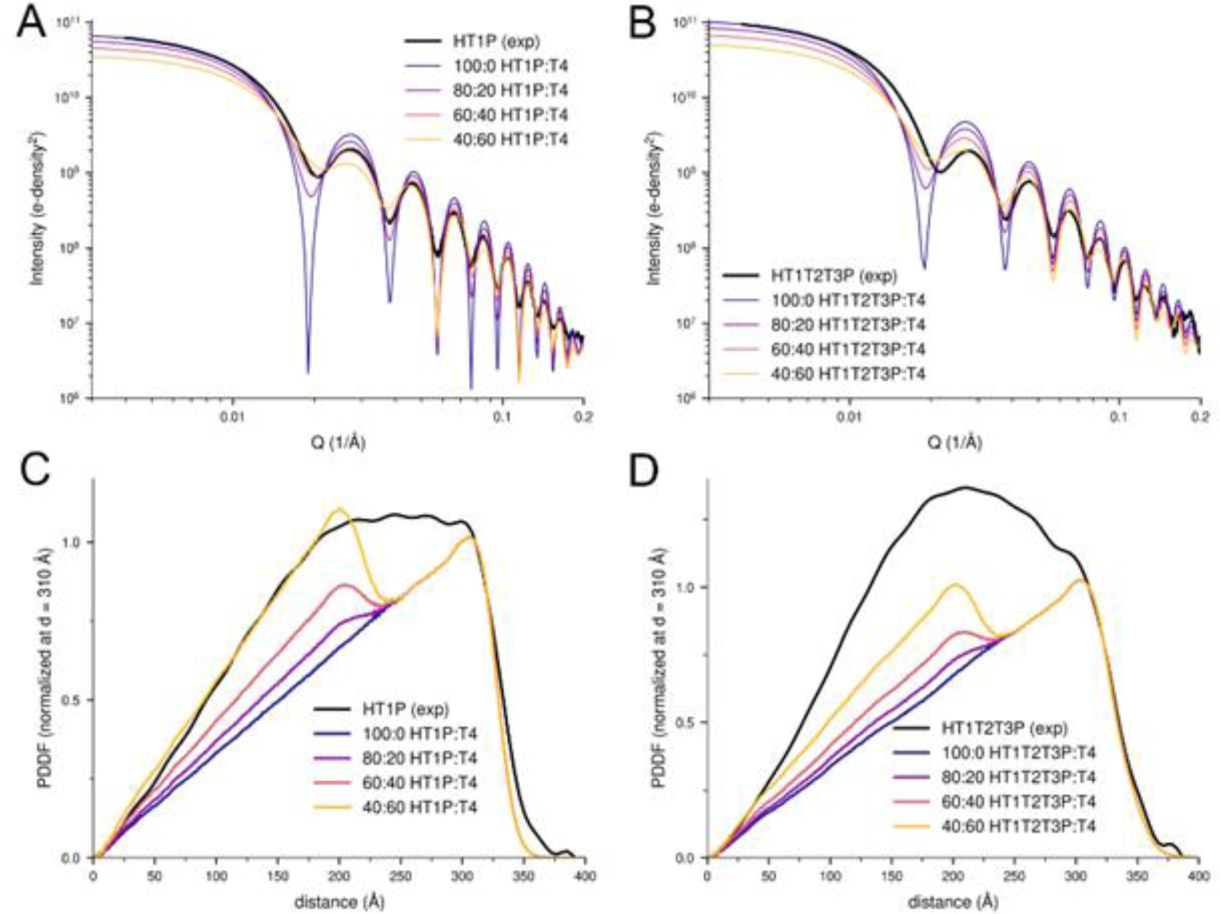
Effect of mixing HTP and HP shells on SAXS and PDDF data. Parts A and B compare experimental (black) and calculated (colored) SAXS profiles for mixtures of HT1P or HT1T2T3P shells with the T4 version of the HP shell, with ratios ranging from 100:0 to 40:60. Parts C and D show the corresponding PDDF profiles, normalized at 310 Å. The addition of HP shells introduces significant variations in SAXS profiles, notably decreasing the depth and shifting the position of the first minimum. However, these variations are still insufficient to explain the PDDF data.

**Figure 8.**
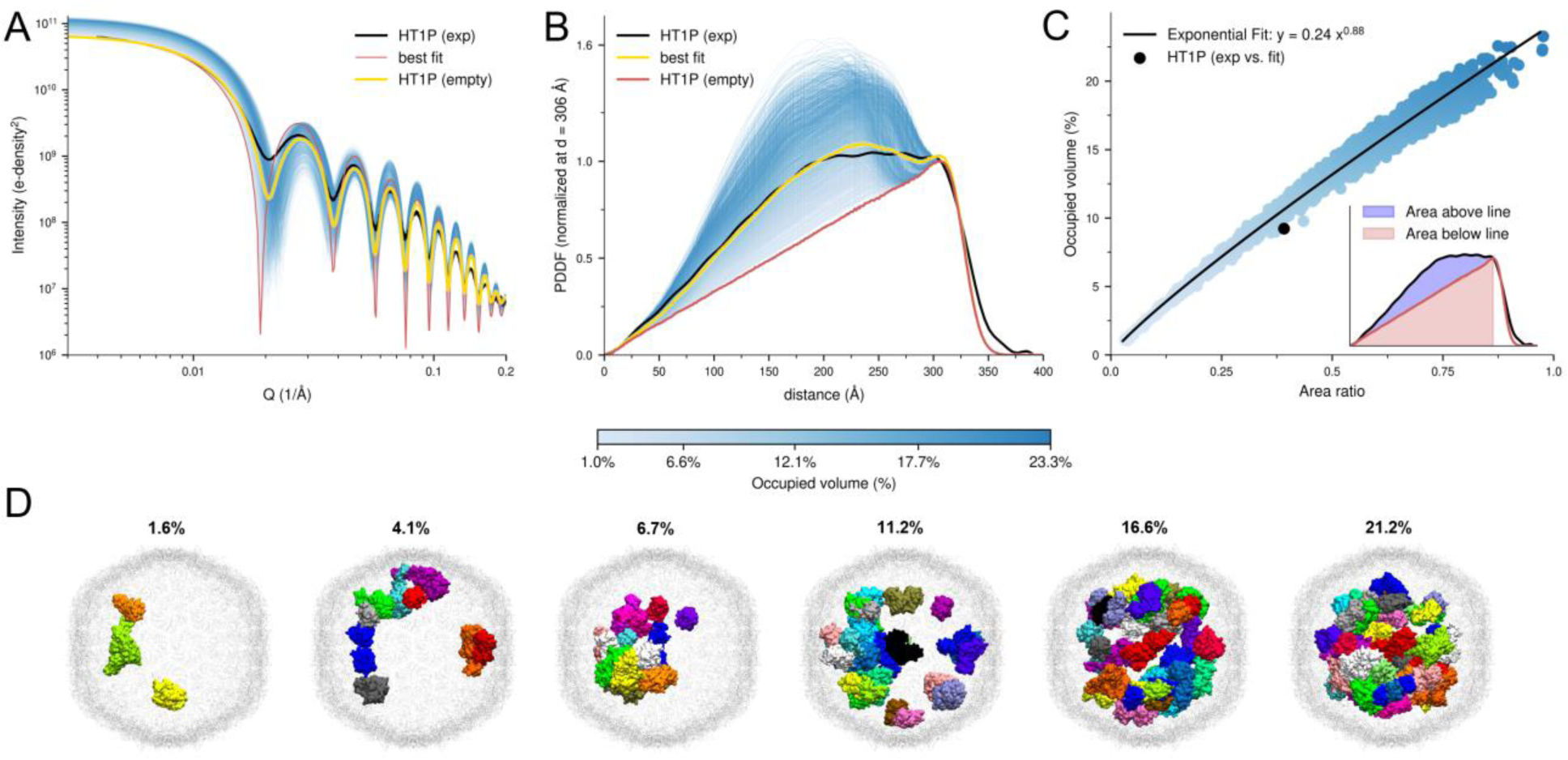
Impact of Cargo Enzymes on SAXS and PDDF Profiles in HT1P Shells. Part A compares the experimental (black) and calculated (colored) SAXS profiles for HT1P shells containing cargo proteins. The blue-to-red colored profiles represent varying cargo-occupied volume fractions, and the yellow line indicates the best fit using an ensemble of 50 structures. Part B shows the corresponding PDDF profiles, normalized at 306 Å. Part C illustrates the strong connection between the cargo-occupied volume fraction and the ratio of the area above and below the line between 0 and 306 Å. This relationship allows the PDDF profile to serve as a direct estimator of the cargo-occupied volume. Part D presents a selection of several cargo-filled shell structures from the fitted ensemble, showing a high diversity in cargo packing within the shell. The values above the structures are the respective cargo-occupied volume.

**Figure 9.**
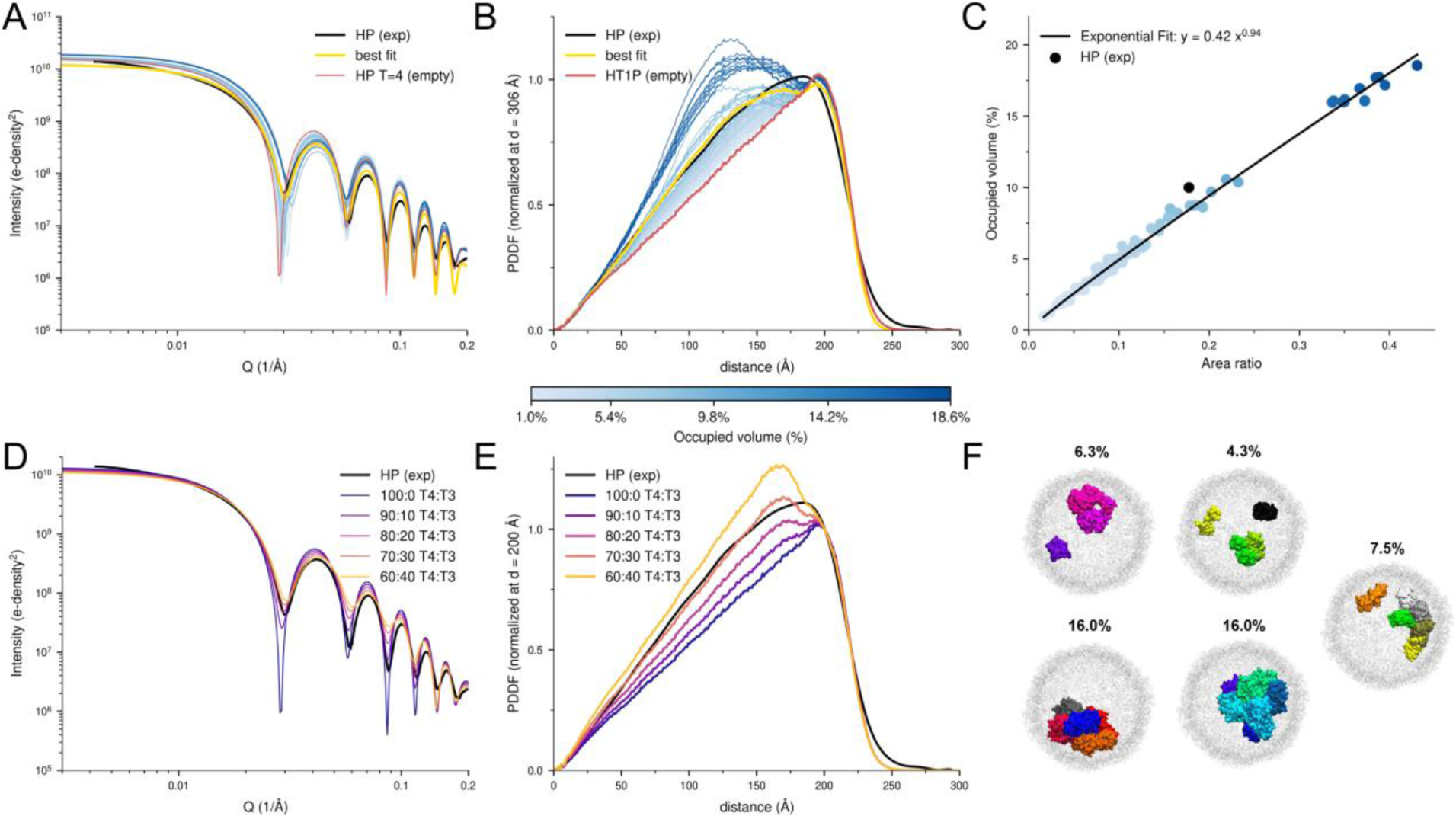
Effect of cargo loading and T=3/T=4 mixtures on SAXS and PDDF profiles of HP shells. Part A compares the experimental (black) and calculated (colored) SAXS profiles for HP shells (T=4) with varying cargo-occupied volume fractions. The blue-to-red gradient represents the different volume fractions occupied by cargo proteins, and the yellow line indicates the best fit using an ensemble of 5 structures. Part B shows the corresponding PDDF profiles, normalized at 200 Å. Part C demonstrates the strong relationship between the cargo-occupied volume fraction and the ratio of the area above and below the line between 0 and 200 Å, allowing direct estimation of the occupied volume from the PDDF profile. Part D compares the experimental (black) and calculated (colored) SAXS profiles for mixtures of HP T=4 and T=3 shells. Part E shows the corresponding PDDF profiles for these mixtures. Part F presents the five cargo-filled HP structures from the fitted ensemble. The values above the structures are the respective cargo-occupied volume. Overall, the results suggest that the HP SAXS profile by itself lacks sufficient information to distinguish between cargo loading and T=3/T=4 shell mixtures.

HO HTP BMC shells exhibit structural heterogeneity, in part due to the presence of three naturally occurring trimer species. The most significant distinction is that T1 trimers are single-layered, while T2 and T3 are double-stacked trimers, Figure 1. Additionally, “wiffle ball” structures without pentamers have been observed,^40^ suggesting the possibility that fully assembled shells might have holes due to missing pentamer tiles.

When we altered the trimer composition in our models, Figures 6A and 6D, we observed minor changes in both the SAXS and PDDF profiles. Increasing the proportion of T3 trimers led to a reduction in the depth of the minima in the SAXS profile and a general increase in intensity due to the larger system size. This was also noted above, in Figure 2, comparing SAXS for HT1P and HT1T2T3P shells. While this affected the absolute values in the PDDF, Figure S8, normalizing the PDDF profiles at a common distance showed that trimer ratio changes resulted similar profiles with the most notable variation seen by the double layered T3 contributions to pair distances above ∼300 Å, Figure 6D. Therefore, variations in trimer composition alone cannot account for the experimental PDDF curves which show significantly higher relative intensities below ∼300 Å for HT1P and HT1T2T3P shells. Similarly, changes in pentamer content had a limited impact on SAXS and PDDF profiles, Figures 6B and 6E, respectively. Reducing pentamer content caused a dampening of the oscillatory SAXS features but only minor changes in the PDDF’s triangular shape.

To explore the role of protein dynamics, we performed an atomistic MD simulation started from the 6mzx cryo-EM structure and calculated the SAXS and PDDF profiles across different frames. Given the large system size (around 10 million atoms for a fully solvated shell), the length of the simulation was limited to 100 ns, sufficient to capture local fluctuations but not large-scale structural changes. During this time frame, only minor damping of the first well in the SAXS spectrum was observed (Figure 6C), leading to small changes in PDDF profiles (Figure 6F). We conclude from this analysis that structural changes in trimer composition, pentamer content, and local dynamics are insufficient to explain the discrepancies between the model and experimental data.

Given that hexamers and pentamers of the HO HTP shells can form smaller HP shells, we examined whether a mixture of HTP (HT1P or HT1T2T3P) shells with the T=4 version of HP shells could account for the discrepancies, Figure 7. The SAXS profiles for modeled mixtures of HTP and HP shells, Figures 7A, B, show that incorporating 20% HP content makes the minima significantly shallower. As the proportion of HP shells increases, we observe a notable shift in the position of the first minimum. For a ∼40% HP content, the position of the first minimum matches the HT1P shell, though this is not the case for HT1T2T3P shells. These simulations show that even by introducing 20% to 40% HP content in HTP shell mixtures, the impact on the PDDF profiles, Figures 7C, D is not sufficient to account for the deviation from the experimental PDDF. The presence of an HP content is seen to increase the pair distance distribution below ∼200 Å but has no effect on the distribution between 200 and 300 Å. As a result, the calculated PDDF profiles still show high deviations from the experimental data. Therefore, a mixture of HP and HTP shells is insufficient to fully explain the discrepancies between the experimental scattering and distance distribution patterns. Further, as discussed above, considering the possibility of purified HT1P and HT1T2T3P shell samples as mixtures with significant mole fractions of multiple shell types can be ruled out from Guinier and TEM^6^ analyses.

As BMCs function to enclose various enzymes, we investigated the influence of encapsulated cargo proteins on the SAXS and PDDF profiles of HT1P shells. Using untargeted proteomics,^34^ we identified cytoplasmic *E. coli* proteins in our samples, with around 10% of the HTP tile abundance (Figure S11). Based on the most abundant proteins from the proteomics data (Table S1), we generated 2400 cargo models with varying amounts and spatial distributions of proteins within the HT1P shell. Figure 8A shows the SAXS profiles of all models, with a blue color gradient indicating different occupied volume fractions of cargo proteins. As the volume fraction increased, the oscillatory features were dampened, and the position of the first minima shifted. The corresponding PDDF profiles, Figure 8B, were similarly affected, with higher cargo volume fractions resulting in increased distance distributions in the 0 to 300 Å range, most prominently between 125 and 275 Å. The PDDF profiles also transitioned from a triangular to a more bell-shaped form.

No single cargo model fully reproduced the experimental SAXS and PDDF profiles. Instead, the average over an ensemble of 50 structures provided the best fit (yellow line). Figure 8D shows a subset of the ensemble, highlighting the diversity in protein number and packing arrangements within the shell. The best-fit ensemble showed significantly better agreement with the experimental PDDF profile, especially in the flatter region between 175 and 310 Å. The ensemble also shifted the first minimum in the SAXS profile to match the experimental minimum and dampened the oscillatory features. These results indicate that while cargo proteins influence the overall scattering and distance distribution, their packing is not uniform. We note that a wide range of potential ensembles with similar volume occupation provides comparable agreement with the experimental data, suggesting flexibility in how cargo can be accommodated within the shell.

Assuming that the primary deviation between experimental PDDF profiles and empty shell models is due to the presence of cargo proteins, we derived a method to directly estimate the occupied volume fraction from experimental PDDF data, Figure 8C. Empty shells form a triangular-shaped PDDF curve, peaking at the average shell diameter. By analyzing the ratio between the area (A_ratio_) above and below the HTP empty shell PDDF from 0 to ∼306 Å, we found that the cargo-occupied volume fraction, V_frac_, can be estimated as follows:

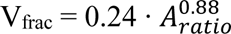

The volume fraction estimated through this formula produces results consistent with the ensemble-averaged values. We applied this method to the 2400 cargo models of HT1T2T3P shells (Figure S12), observing a similar cargo effect as seen in the HT1P shell. For the HT1T2T3P shell, a cargo-based ensemble achieved high agreement with the experimental SAXS and PDDF profiles. The key difference, however, is that the estimated volume fraction was approximately 16%, nearly double the volume fraction found for the HT1P shell (∼9%). For the HP shell, Figure 9, combining cargo-based ensembles with a mixture of T=4 and T=3 shells produced a comparable level of agreement with the experimental data. This was likely due to the relatively small size differences between the T=3 and T=4 HP shells, making the shell mixing and cargo-loading effects harder to distinguish in the SAXS and PDDF profiles.

## DISCUSSION

Both ab initio and molecular modeling point toward the finding that adventitious capture of *E. coli* cytoplasmic proteins during BMC shell assembly is the most plausible explanation for the deviations between experimental SAXS and PDDF profiles and those calculated from empty shell models. Molecular modeling demonstrated that changes in shell composition, such as variations in trimer ratios or missing pentamers, as well as local fluctuations, had only a minimal effect on the SAXS and PDDF profiles. While mixtures of shell structures and sizes were found to impact SAXS profiles by dampening the oscillatory patterns, quantities of contaminating shell structures compatible with SAXS and TEM analyses of homogeneity were found to have limited influence on PDDF profiles. In contrast, the presence of plausible amounts of cytoplasmic proteins was found to substantially influence both SAXS and PDDF profiles by reducing the overall size, shifting the position of the minima, and increasing the distance distribution in the critical mid-range region in accord with experimental data. This highlights the significant role cargo plays in shaping the experimental scattering patterns, making it the key factor in explaining the observed deviations.

Having identified that deviations between experimental and calculated PDDFs for purified BMC shell samples can be assigned to <u>adventitious</u> capture of cytoplasmic cargo, it is possible to estimate the quantity of cargo captured during shell assembly directly from the difference of the integrated PDDFs because the square root of the PDDF area is proportional to the total electrons or the molecular weight (MW) of the molecular assembly.^36-38^ For example, from the experimental PDDFs for HT1P, HT1T2T3P, and HP shells that are scaled to those of reference structures in Figure 2C, 2D, 3B, we obtain the area ratios between experiment and reference PDDFs of 1.37, 1.56 and 1.25, respectively. From the square-rooted area ratios and the reference theoretical MWs, we obtain cargo protein per shell that are 0.9 MDa, 1.5 MDa and 0.2 MDa for HT1P, HT1T2T3P and HP, respectively. However, adventitious cargo loading seems random and varies among preparations. For example, estimates of the extent of adventitious cargo loading in *E. coli* expressed HT1P shells varied from 0.18 MDa to 0.46 MDa in three preparations presented in Figure S10B.

That BMC shells without recognition tags adventitiously capture significant amounts of cytoplasmic cargo has not been previously recognized. Understanding this phenomenon could be critical for the design of BMC shells that target selected enzyme capture. Given the extremely restricted volume within nanoscale BMC shells, the presence of adventitious cargo would compete with enzyme targets for a limited amount of interior space. For example, in shells with recognition tags for selected enzyme encapsulation, SDS-PAGE, immunoblot analysis, and mass spectroscopy have demonstrated elevated presence of targeted enzymes in purified shell samples, although TEM and cryo-EM have had only limited success in identifying the location of these captured proteins.^5, 16, 35, 36^ In the case of cryo-EM measurements, variability in the images showing electron density in the shell interior prevented image indexing and construction of maps for captured enzyme localization.^5^ We note that the nanoscale BMC shells are below the resolution limits for fluorescence imaging microscopy. Similarly, while dynamic light scattering is widely used for BMC shell characterization, it is sensitive to the shell size but not interior content.

Hence, we find the presented method of comparison of experimental PDDF to reference patterns calculated from reference structures provides a possibly unique opportunity to characterize the extent of protein cargo captured during BMC shell assembly. One of the advantages of this approach is that SAXS measurements are a high-throughput assay that can be directly applied to the BMC shell solutions used for biochemical enzyme activity assays. We expect that this approach will prove to be valuable for the development and evaluation of BMC shell architectures for biological and abiotic catalyst capture and as platforms for constructing compartments for catalysis in confinement.

## ACKNOWLEDGMENT

This work was supported as part of the Center for Catalysis in Biomimetic Confinement, an Energy Frontier Research Center funded by the U.S. Department of Energy, Office of Science, Basic Energy Sciences. At Michigan State University the work was supported under award DE-SC0023395, at Argonne National Laboratory under contract DEAC02-06CH11357, and at Lawrence Berkeley National Laboratory under contract DE-AC02-05CH11231. This research used resources of the Advanced Photon Source operated by Argonne National Laboratory under Contract DEAC02-06CH11357 and the National Synchrotron Light Source II operated by Brookhaven National Laboratory under Contract DE-SC0012704, both US DOE Office of Science user facilities.

