## Supporting Information for "Structure Characterization of Bacterial Microcompartment Shells via X-ray Scattering and Coordinate Modeling: Evidence for adventitious capture of cytoplasmic proteins"

[1]X-ray Science Division, Argonne National Laboratory, Lemont, IL, 60439 USA, [2]Department of Biochemistry & Molecular Biology, Michigan State University, East Lansing, MI 48824 USA, [3]Chemical Sciences and Engineering Division, Argonne National Laboratory, Lemont, IL, 60439 USA, [4]MSU-DOE Plant Research Laboratory, Michigan State University, East Lansing, MI, 48824 USA, [5]Molecular Foundry Division, Lawrence Berkeley National Laboratory, Berkeley, CA, 94720 USA, [6]Molecular Biophysics and Integrated Bioimaging Division, Lawrence Berkeley National Laboratory, Berkeley, CA, 94720 USA, [7]Environmental Genomics and Systems Biology Division, Lawrence Berkeley National Laboratory; Berkeley, CA, 94720 USA.

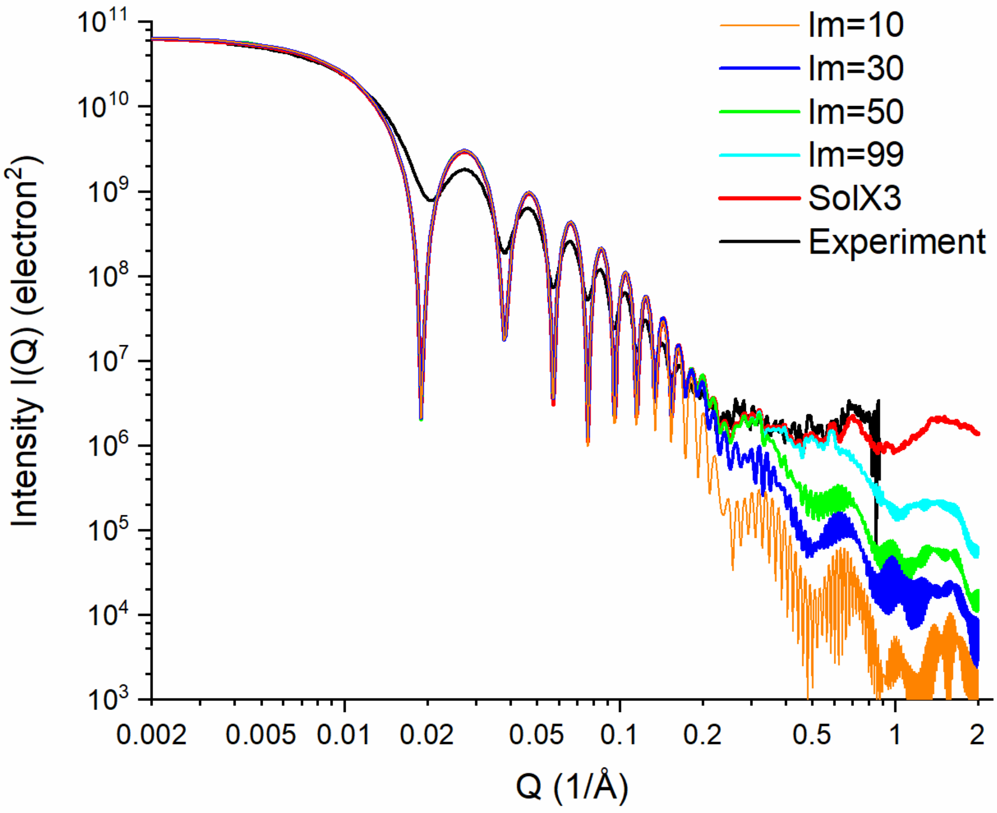


Figure S1. Comparison of SAXS using spherical harmonics expansion in CRYSOL (with various levels, lm=10, 30, 50, 99), atomic form factors as implemented in SolX3 (red line), and experimental data (black line).

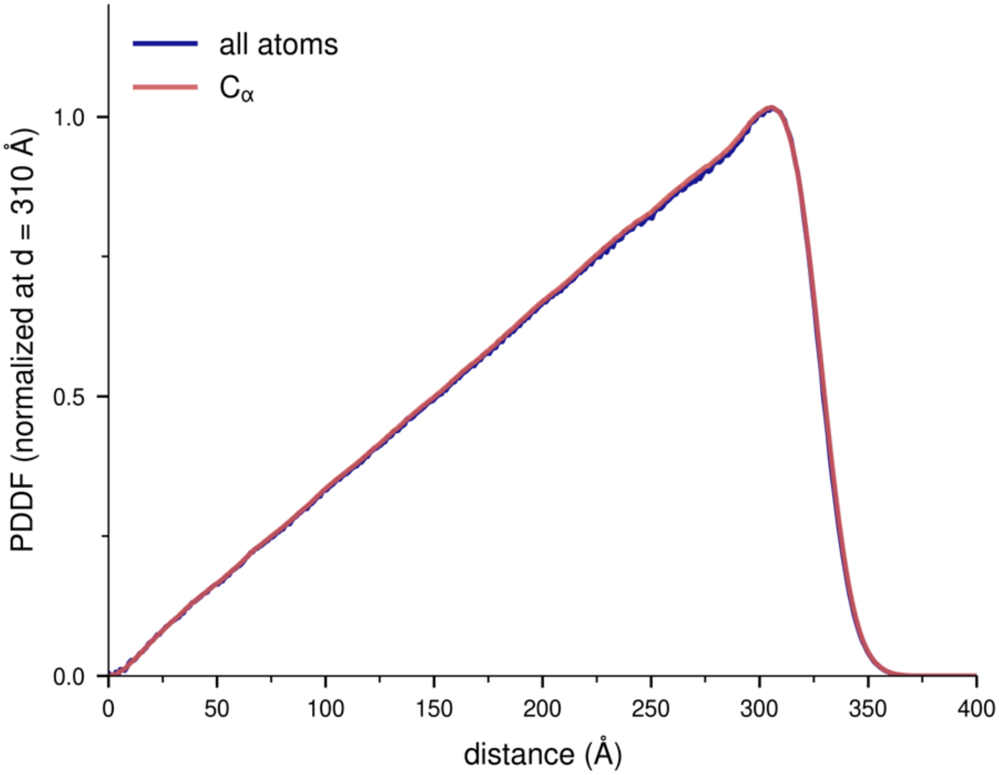


Figure S2. PDDF profiles calculated for a HT1P shell using all atoms (blue line) versus only $C_\alpha$ (red line) atoms for the PDDF calculation.

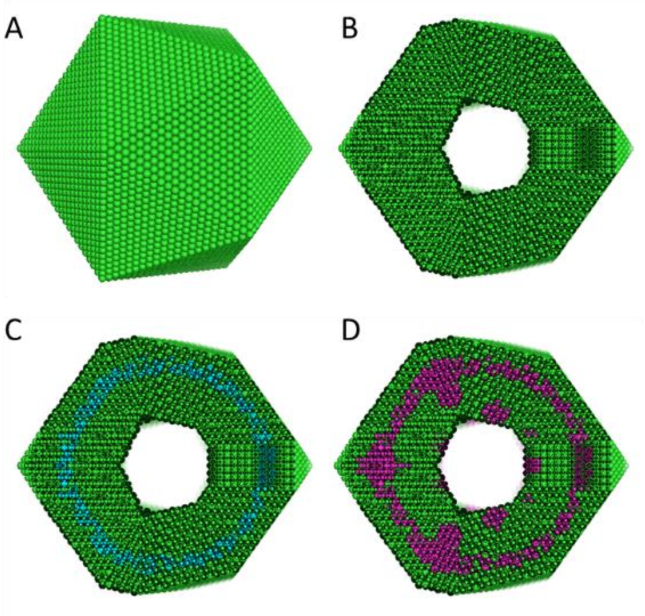


Figure S3. Search space volumes used for DMMIN calculations. Overall (A) and cross-section (B) views of the search space of DAMMIN reconstruction for HT1P reference shell structure. This search space was used for fitting SAXS both calculated from the HT1P coordinates and from experimental data. The search space is composed of densely packed small beads of radius 5.7 Å. C & D: Cross-section views of reconstructed structures within the search volume. (C) Cyan beads mark the structure re-constructed from the calculated HT1P SAXS profile. (D) Magenta beads mark the structure fit to experimental data. The structures are the exact ones are also shown in Fig 5.

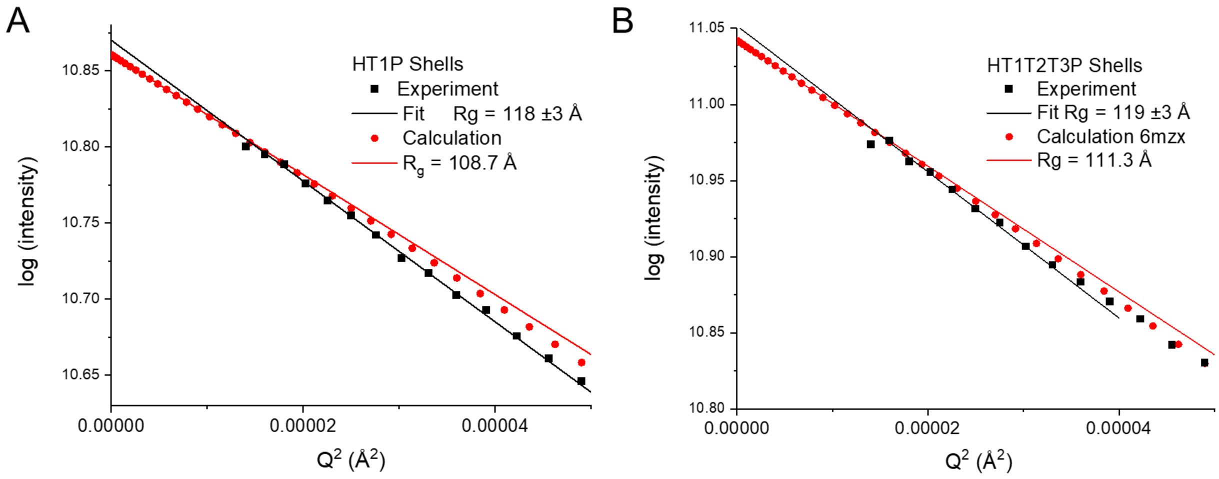


Figure S4. Guinier plots for HT1P (Part A) and HT1T2T3P (Part B) shells. Experimental data and calculation from the respective reference structures are marked by black and red symbols, respectively. The solid lines are fits using the Guinier expression: $\log[I(q)] = I(0) - (R_g q)^2/3$. Values $R_g$ determined from the fitting are shown in the inserts.

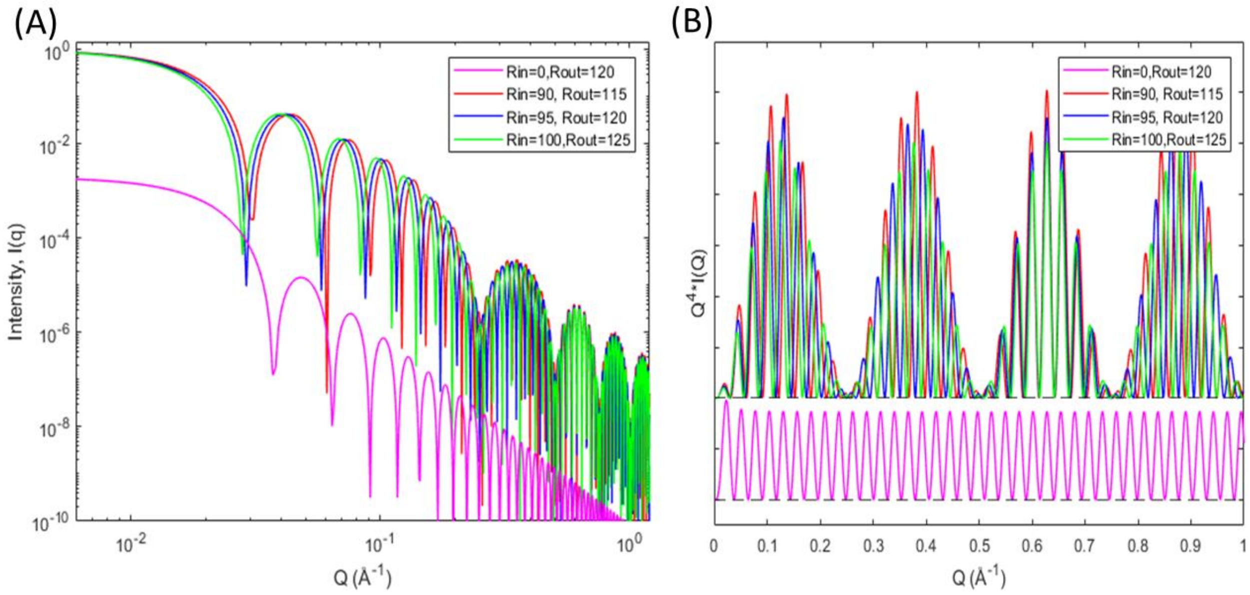


Figure S5. X-ray scattering features of solid sphere, and hollow core-shell spheres. (A). Normalized scattering profiles for hollow spheres. The solid sphere scattering profile was vertically offset for clarity. (B). Same profiles in (A) but plotted in $Q^4*I(Q)$ vs Q presentation, also known as Porod-Debye plot. Solid sphere profile was offset for clarity. Dash lines are zero lines. Magenta: solid sphere, outer radius (Rout) = 125 Å; Red: hollow core-shell sphere, inner radius (Rin) = 90 Å and outer radius = 115 Å; Blue: hollow core-shell sphere, Rin=95 Å, Rout=120 Å; Green: hollow core-shell sphere, Rin=100 Å, Rout=125 Å. All three hollow spheres have same shell wall thickness of 25 Å. In these geometric models, the electron density was assumed to be uniform in the solid portion.

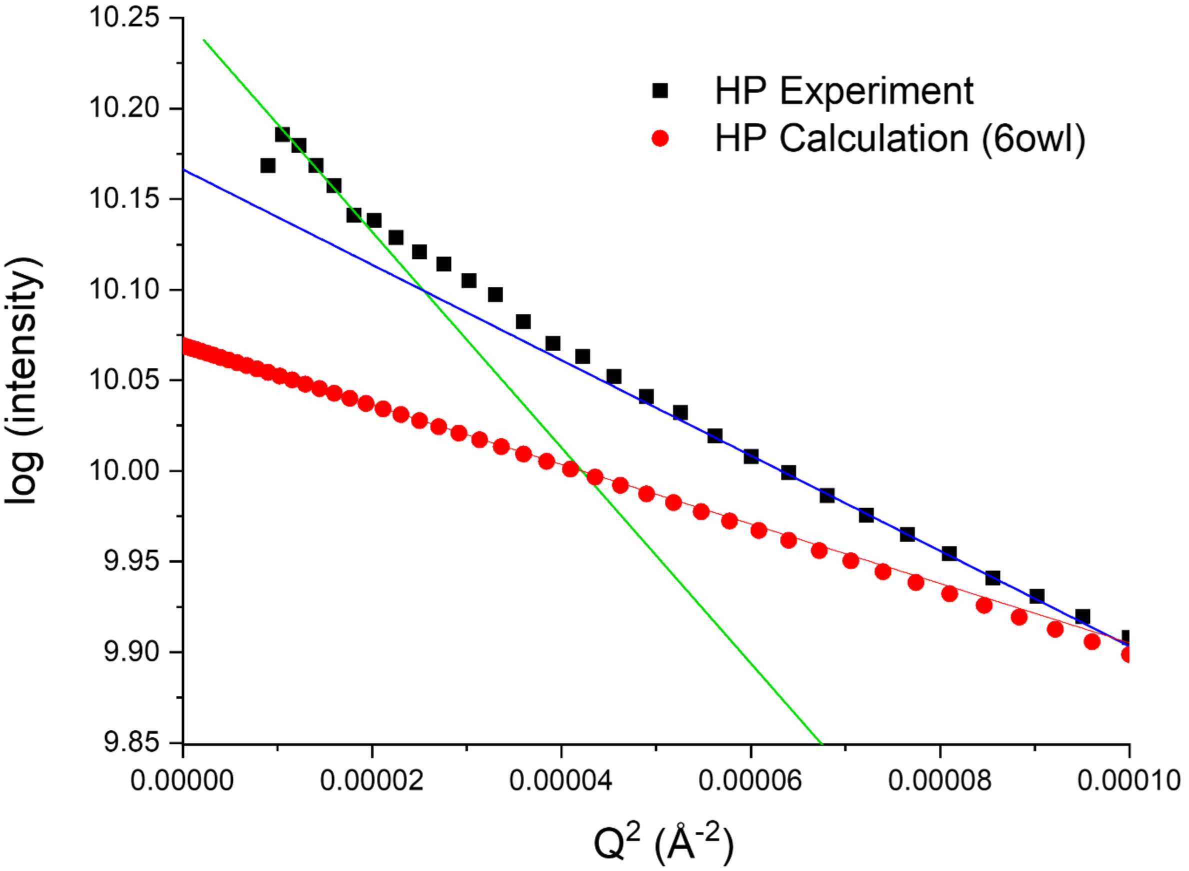


Figure S6. Guinier plots for experimental (black squares) and calculated (red dots) SAXS for HP shells. The green line shows a Guinier fit to low angle experimental scattering, corresponding to an $R_s$ of 134 ±5 Å; the blue is a Guinier fit to higher angle experimental scattering data, corresponding to an $R_s$ of 89 ±0.8 Å. The scattering calculated from the 6owl structure corresponds to an $R_g$ of 70.7 Å.

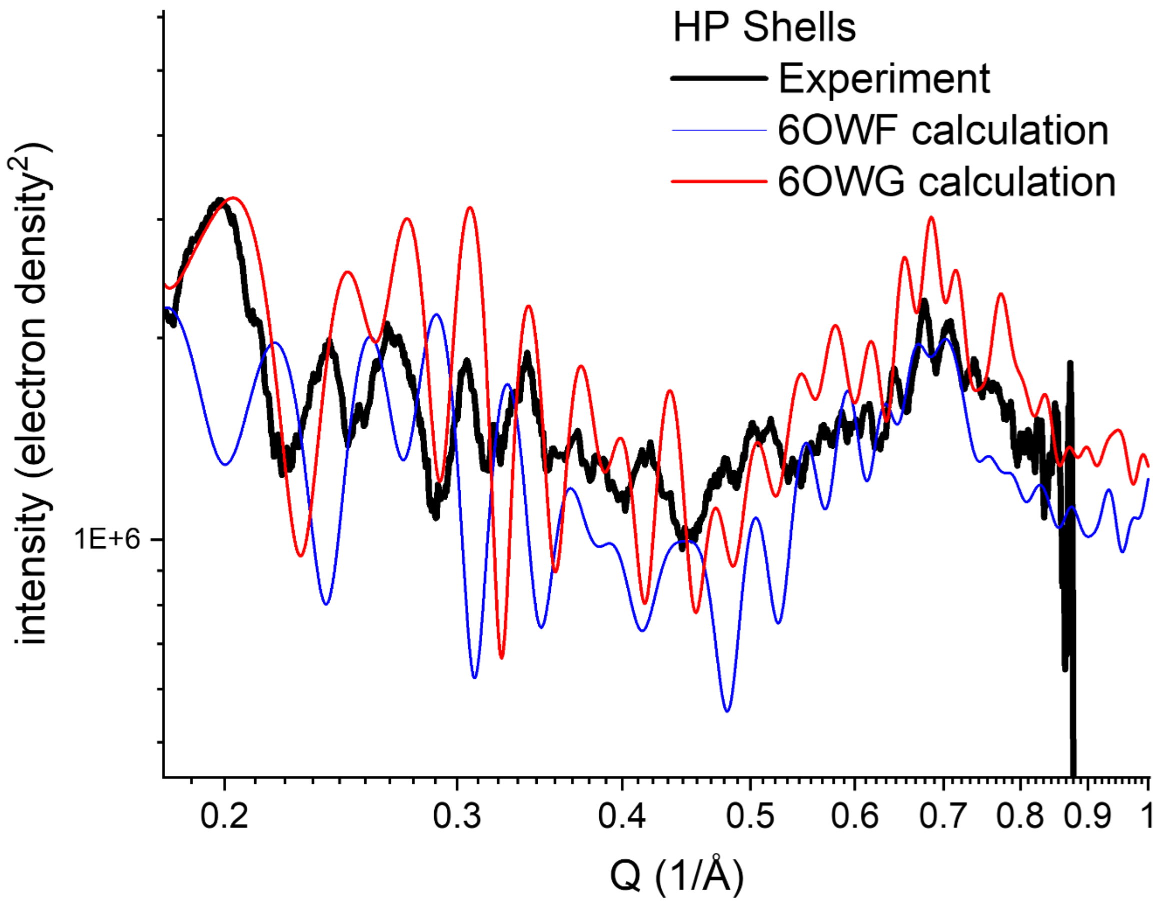


Figure S7. Comparison of wide angle scattering for HO-HP shells measured experimentally (black) with scattering patterns calculated from the T=3 shell, 6owf (blue), and the T=4 shell 6owg (red). The experimental, black, and 6owg calculated, red, curves show approximate correlation in interference frequency patterns.

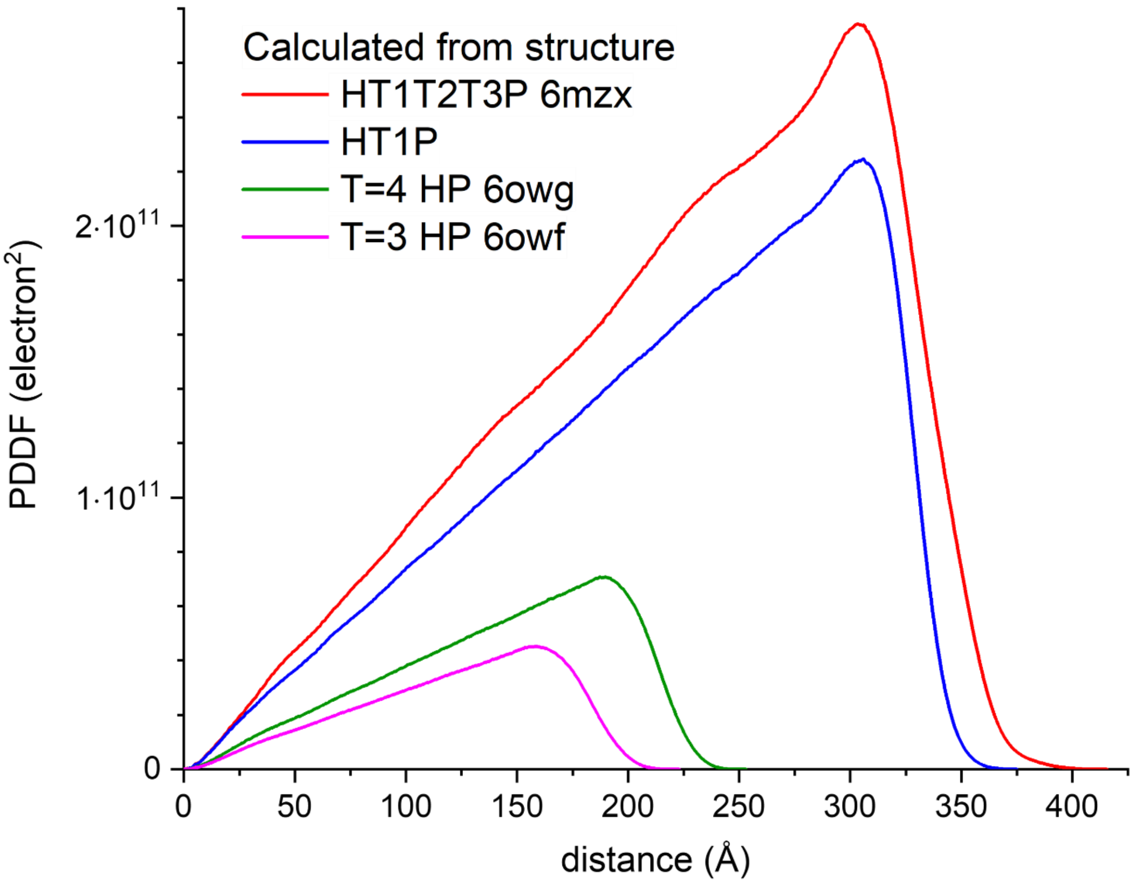


Figure S8. Comparison of PDDF patterns calculated from the BMC shell reference structures.

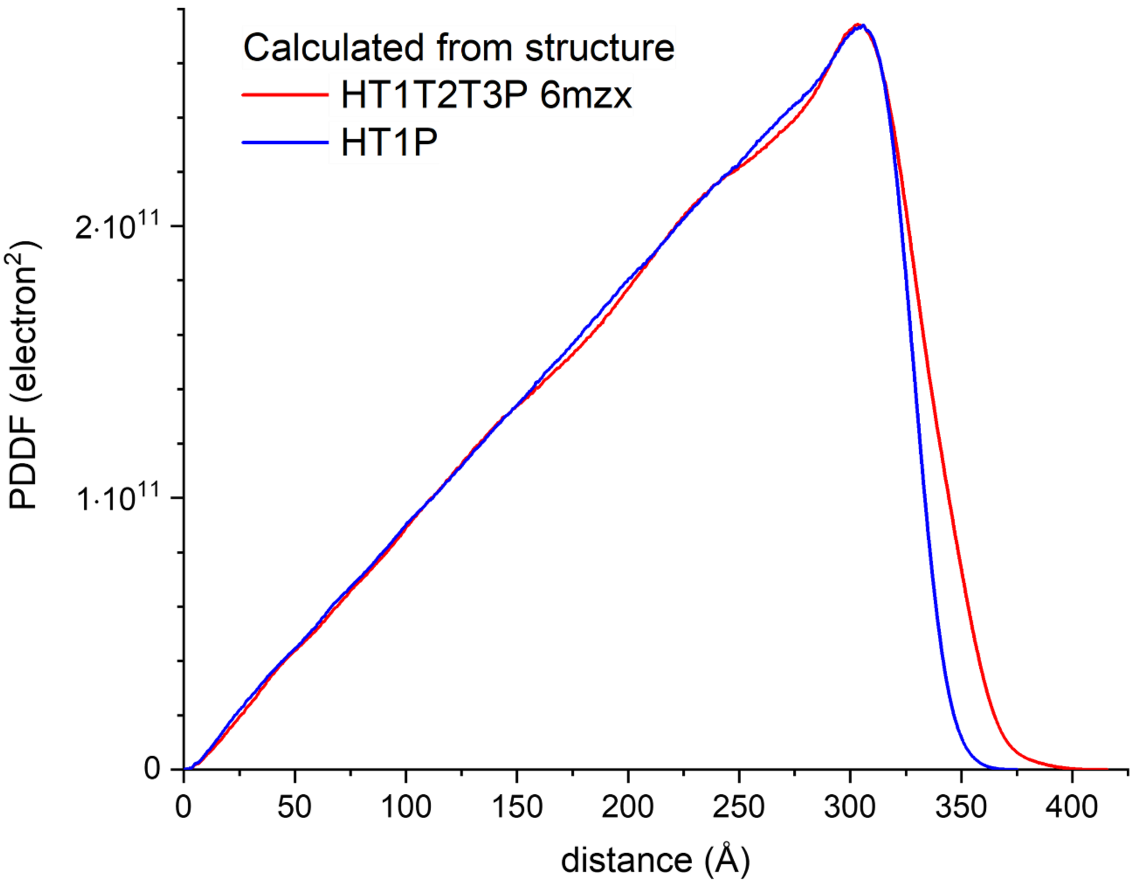


Figure S9. Comparison of PDDF calculated from the HT1T2T3P (red) and HT1P (blue) reference structures. The PDDF for HT1P was scaled to match that of HT1T2T3P.

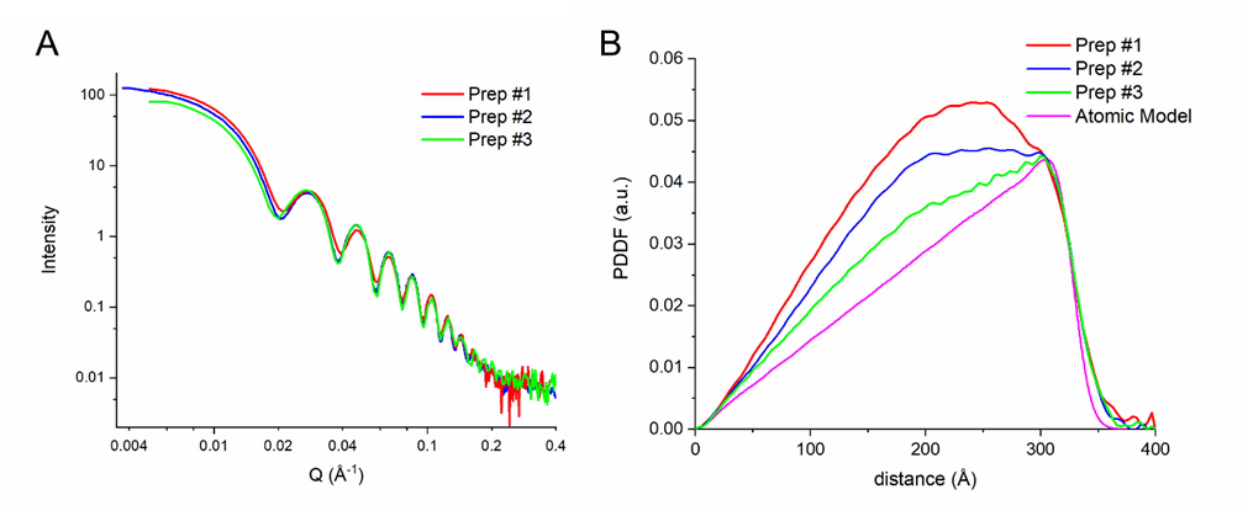


Figure S10. X-ray scattering data (Part A) and PDDFs (Part B) measured from different preparation of *E. coli* expressed HO-HT1P shells.

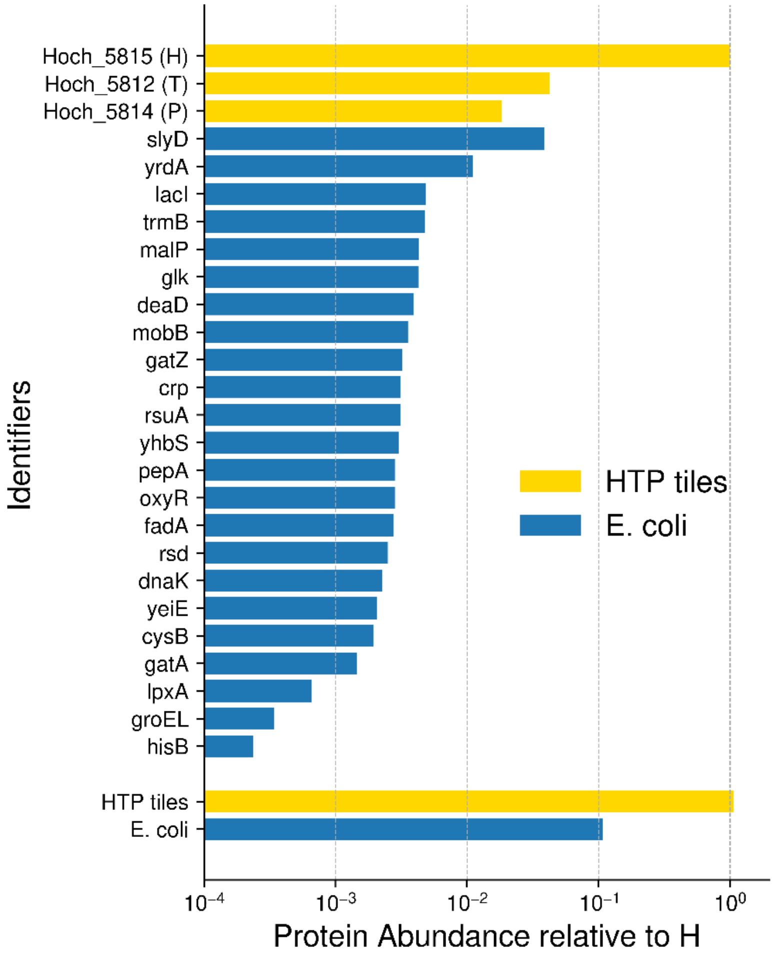


Figure S11. Untargeted proteomic analysis of purified HTP shells. Protein abundances were normalized to the signal of the hexamer subunit to ensure consistency in comparisons. Proteins with relative abundances below $10^{-4}$ were excluded from the analysis, along with common human cell contaminants.

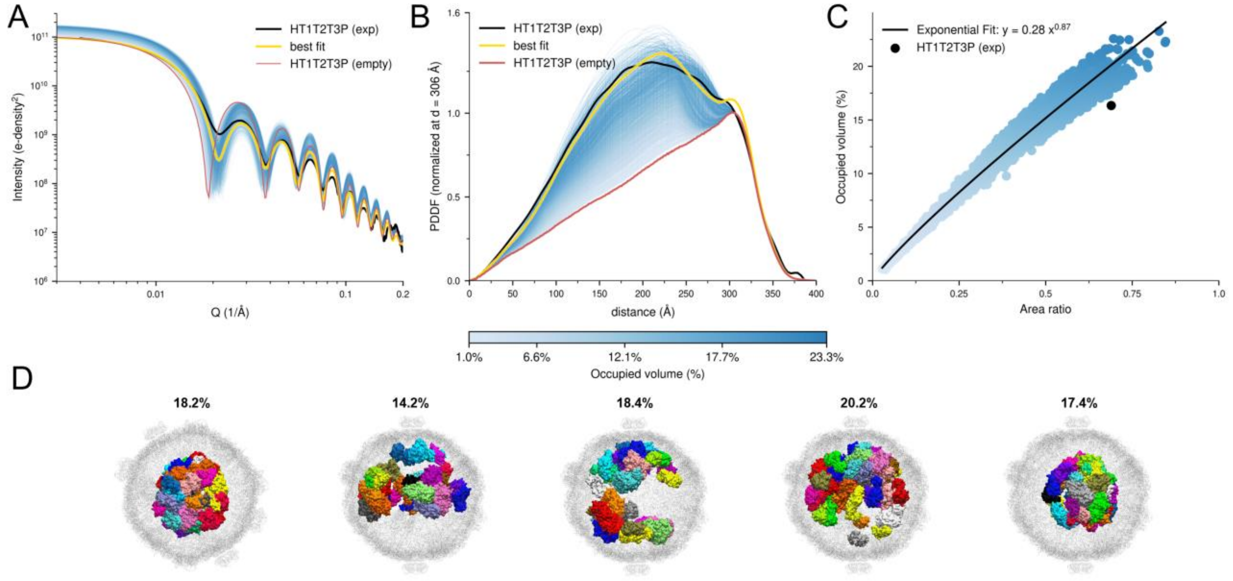


Figure S12. Impact of cargo enzymes on SAXS and PDDF profiles in HT1T2T3P shells. Part A compares experimental (black) and calculated (colored) SAXS profiles for HT1T2T3P shells containing cargo proteins. The blue-to-red colored profiles represent varying cargo-occupied volume fractions, and the yellow line indicates the best fit using an ensemble of 50 structures. Part B shows the corresponding PDDF profiles, normalized at 306 Å. Part C illustrates the strong connection between the cargo-occupied volume fraction and the ratio of the area above and below the line between 0 and 306 Å. This relationship allows the PDDF profile to serve as a direct estimator of the cargo-occupied volume. Part D presents a selection of several cargo-filled shell structures from the fitted ensemble, showing a high diversity in cargo packing within the shell. The values above the structures are the respective cargo-occupied volume.

Table S1: Summary of the untargeted proteomic analysis of purified HTP shells. Proteins with an abundance (normalized total precursor intensity) below $10^{10}$ were excluded, along with common human cell contaminants. The final columns indicate the oligomerization state of the identified proteins used in the modelling of cargo-loaded shells and the origin of the structural model (AF2: AlphaFold2).

| Accession Number | Alternate ID | Molecular Weight | Protein Abundance | Species | Modeled oligomerization | Structure |
|---|---|---|---|---|---|---|
| D0LID5 | Hoch_5815 | 10 kDa | 5.19E+13 | *HO* | Hexamer | |
| D0LHE3 | Hoch_5812 | 22 kDa | 2.19E+12 | *HO* | Trimer | |
| D0LHE5 | Hoch_5814 | 10 kDa | 9.59E+11 | *HO* | Pentamer | |
| P0A9K9 | slyD | 21 kDa | 2.01E+12 | *E. coli* | Monomer | 2KFW |
| P0A9W9 | yrdA | 20 kDa | 5.75E+11 | *E. coli* | Trimer | 3TIO |
| P03023 | lacI | 39 kDa | 2.51E+11 | *E. coli* | Dimer | 1EFA |
| P0A8I5 | trmB | 27 kDa | 2.48E+11 | *E. coli* | Monomer | 3DXX |
| P00490 | malP | 91 kDa | 2.23E+11 | *E. coli* | Dimer | 1AHP |
| B1X9R0 (+2) | glk | 35 kDa | 2.22E+11 | *E. coli* | Monomer | AF2 |
| P0A9P6 | deaD | 71 kDa | 2.03E+11 | *E. coli* | Dimer | AF2 |
| P32125 | mobB | 19 kDa | 1.85E+11 | *E. coli* | Dimer | 1NP6 |
| C4ZSH9 (+1) | gatZ | 47 kDa | 1.68E+11 | *E. coli* | Monomer | AF2 |
| P0ACJ8 | crp | 24 kDa | 1.63E+11 | *E. coli* | Dimer | 1G6N |
| P0AA43 | rsuA | 26 kDa | 1.61E+11 | *E. coli* | Monomer | 1KSK |
| P63417 | yhbS | 19 kDa | 1.57E+11 | *E. coli* | Monomer | AF2 |
| B1XEN9 (+2) | pepA | 55 kDa | 1.47E+11 | *E. coli* | Hexamer | 1GYT |
| P0ACQ4 | oxyR | 34 kDa | 1.47E+11 | *E. coli* | Monomer | AF2/1I6A |
| P21151 | fadA | 41 kDa | 1.44E+11 | *E. coli* | Monomer | AF2 |
| B1XBZ8 (+2) | rsd | 18 kDa | 1.30E+11 | *E. coli* | Monomer | 4XWJ |
| P0A6Y8 | dnaK | 69 kDa | 1.17E+11 | *E. coli* | Monomer | 4B9Q |
| P0ACR4 | yeiE | 33 kDa | 1.07E+11 | *E. coli* | Monomer | AF2 |
| P0A9F3 | cysB | 36 kDa | 1.01E+11 | *E. coli* | Monomer | AF2 |
| P69828 | gatA | 17 kDa | 7.54E+10 | *E. coli* | Monomer | AF2 |

| | | | | | | |
|---|---|---|---|---|---|---|
| B1XD50 (+2) | lpxA | 28 kDa | 3.42E+10 | *E. coli* | Trimer | 1LXA |
| B1XDP7 (+2) | groEL | 57 kDa | 1.78E+10 | *E. coli* | -* | -* |
| P06987 | hisB | 40 kDa | 1.23E+10 | *E. coli* | Monomer | AF2 |

*did not model groEL due to large size
